# Myelin speeds cortical oscillations by consolidating phasic parvalbumin-mediated inhibition

**DOI:** 10.1101/2021.09.07.459122

**Authors:** Mohit Dubey, Maria Pascual-Garcia, Koke Helmes, Dennis D. Wever, Mustafa S. Hamada, Steven A. Kushner, Maarten H. P. Kole

## Abstract

Parvalbumin-positive (PV^+^) γ-aminobutyric acid (GABA) interneurons are critically involved in producing rapid network oscillations and cortical microcircuit computations but the significance of PV^+^ axon myelination to the temporal features of inhibition remains elusive. Here using toxic and genetic models of demyelination and dysmyelination, respectively, we find that loss of compact myelin reduces PV^+^ interneuron presynaptic terminals, increases failures and the weak phasic inhibition of pyramidal neurons abolishes optogenetically driven gamma oscillations *in vivo*. Strikingly, during periods of quiet wakefulness selectively theta rhythms are amplified and accompanied by highly synchronized interictal epileptic discharges. In support of a causal role of impaired PV-mediated inhibition, optogenetic activation of myelin-deficient PV^+^ interneurons attenuated the power of slow theta rhythms and limited interictal spike occurrence. Thus, myelination of PV axons is required to consolidate fast inhibition of pyramidal neurons and enable behavioral state-dependent modulation of local circuit synchronization.

## Introduction

GABAergic interneurons play fundamental roles in controlling rhythmic activity patterns and the computational features of cortical circuits. Nearly half of the interneuron population in the neocortex is parvalbumin-positive (PV^+^) and comprised mostly of the basket-cell (BC) type ^1, 2^. PV^+^ BCs are strongly and reciprocally connected with pyramidal neurons (PNs) and other interneurons, producing temporally precise and fast inhibition ^3–5^. The computational operations of PV^+^ BCs, increasing gain control, sharpness of orientation selectively and feature selection in the sensory cortex ^6–10^ are mediated by a range of unique molecular and cellular specializations. Their extensive axon collaterals targeting hundreds of PNs, are anatomically arranged around the soma and dendrites, electrotonically close to the axonal output site and the unique calcium (Ca^2+^) sensors in PV^+^ BCs terminals, synaptotagmin 2 (Syt2), are tightly coupled to Ca^2+^ channels mediating fast and synchronized release kinetics ^11, 12^, powerfully shunting excitatory inputs and increasing the temporal precision of spike output ^2, 5, 13, 14^.

Recent findings have shown that the proximal axons of PV^+^ interneurons are covered by myelin sheaths ^5, 13, 15–18^. How interneuron myelination defines cortical inhibitory interneuron functions remains, however, still poorly understood. Myelination of axons provides critical support for long-range signaling by reducing the local capacitance, producing saltatory conduction and by maintaining the axonal metabolic integrity ^19, 20^. For PV^+^ BCs, however, the average path length between the axon initial segment (AIS) and release sites involved in local circuit inhibition is typically less than ∼200 µm ^5, 21, 22^ and theoretical and experimental studies indicate the acceleration by myelin may play only a limited role in tuning inhibition ^16, 22^. Another notable long-standing hypothesis is that myelination of PV^+^ axons may be critical for the security and synchronous invasion of presynaptic terminals ^13^. In support of a role in reliability, in Purkinje cell axons of the long Evans shaker (*les*) rat, which carries a deletion of *Mbp*, spike propagation shows failures and presynaptic terminals are disrupted ^23^. Interestingly, in a genetic model in which oligodendrocyte precursor cells lack the γ2 GABA_A_ receptor subunit, fast-spiking interneuron axons in the neocortex become aberrantly hypermyelinated and proper feedforward inhibition is also impaired^24^. At the network level PV^+^ BC-mediated feedback and feedforward inhibition produces local synchronization of PN and interneuron firing at the gamma (γ) frequency bandwidth which is a key rhythm binding information from cell assemblies, allowing synaptic plasticity and higher cognitive processing of sensory information ^2, 8, 25–27^. Here we directly addressed whether PV^+^ BC driven cortical rhythms require myelination by longitudinally examining the frequency spectrum of cortical oscillations in de- and dysmyelination models and studying the properties and role of genetically labelled PV^+^ BCs.

## Results

### Behavioral state-dependent increase in theta power and interictal epileptiform discharges

We investigated the cortical rhythms by recording *in vivo* local field potential (LFP) in layer 5 (L5) together with the electrocorticogram (ECoG) from both primary somatosensory (S1) and visual (V1) areas. Freely moving mice (C57BL/6) were recorded during home cage explorative behaviors every second week (18–24 hours/week) across an 8-week cuprizone treatment, inducing toxic loss of oligodendrocytes in white- and gray matter including the sensory cortex ^28–30^. Remarkably, after 6 weeks of cuprizone feeding we detected high-voltage spike discharges (∼5 times the baseline voltage and ∼50–300 ms in duration, **Fig. 1a-c**). These brief spike episodes on the ECoG and LFP (**Fig. 1c**) occurred bilaterally and near synchronously in S1 and V1, resembling the interictal epileptiform discharges (also termed interictal spikes) that are a hallmark of epilepsy ^31–34^. Automated detection of interictal spikes in the raw ECoG–LFP signal was performed with a machine-based learning classifier (see Supplementary **Fig. S1a** and **Methods**), revealing a progressively increasing interictal spike rate from ∼5/hour at 4 weeks up to ∼70/hour at 8 weeks (**Fig. 1d**). Importantly, interictal spikes were highly dependent on vigilance state and present exclusively during quiet wakefulness (30 out of 30 randomly selected LFP segments from awake or quiet wakefulness, Chi-squared test *P* < 0.0001, *n* = 6 cuprizone mice), with no other discernible association to specific behaviors (Supplementary **Fig. S1b** and **Movie S1**). Whether the aberrant cortical oscillations were specific to certain frequency bands, including gamma (γ, 30-80 Hz), was examined by plotting the power spectrum density of the LFP in S1 during periods of quiet wakefulness or active movement (**Fig. 1e**). In cuprizone-treated mice LFP power was selectively amplified during quiet wakefulness in the theta frequency band (θ, 4-12 Hz, **Fig. 1e-f**, and Supplementary **Fig. S1c**). In contrast, during active states when mice were moving and exploring no differences were observed in the power spectrum, in none of the frequency bands (**Fig. 1e, f**). Finally, to examine whether interictal epileptiform discharges are due to off-target effects of cuprizone treatment the dysmyelinated *shiverer* mice (*Mbp*^Shi^), lacking compact myelin due to a truncating mutation in *Mbp* ^35^ were recorded at the age of 8 weeks. *Shiverer* mice suffer progressively increasing number of epileptic seizures beginning at approximately 8 weeks of age ^35, 36^ and while ictal discharges were observed at that age, the mice also exhibited interictal spikes with a rate of ∼60/hour, comparable to cuprizone-treated mice but with substantially longer duration (∼100–500 ms, **Fig. 1g-h** and **Fig. S1d**).

**Fig. 1.**
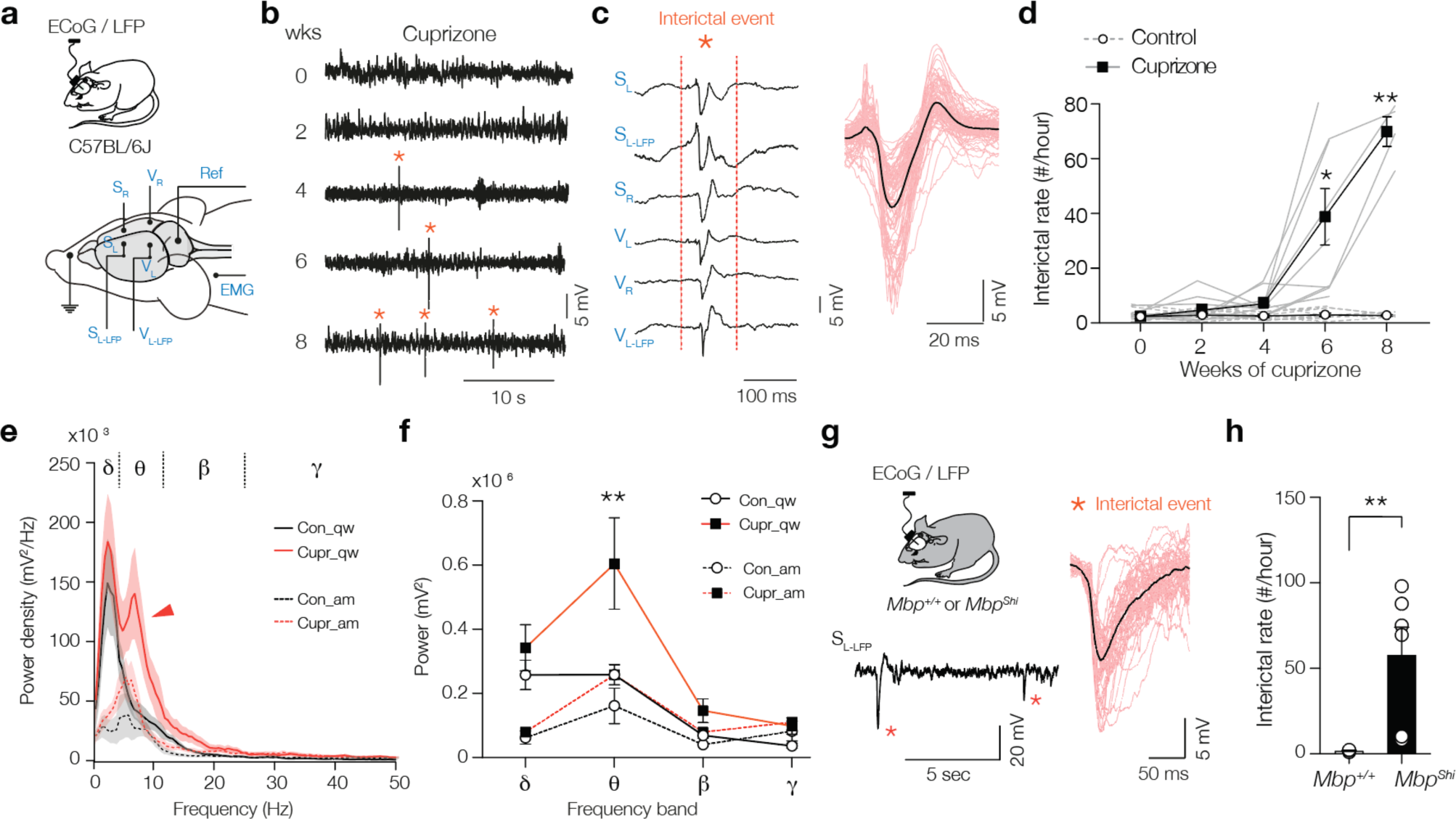
Loss of compact myelin causes interictal spikes and behavioral state-dependent amplification of theta rhythms. (**A**) Schematic of the ECoG and LFP recordings in freely-moving mice. Electrodes were placed right (S_R_) and left (S_L_) in the primary somatosensory cortex, and a left LFP electrode (S_L-LFP_) into L5. A similar array of electrodes was positioned in the primary visual cortex (V_R_, V_L_ and V_L-LFP_). One electrode was placed around neck muscle recording electromyography (EMG) and one used as reference (Ref). (**B**) Interictal spikes (*) appear from 4 weeks cuprizone and onwards. Example raw LFP traces (S_L-LFP_). (**C**) Representative interictal spike example showing spatiotemporal synchronization of the spike across cortical areas and hemispheres. Higher magnification of interictal spikes (red, ∼50 to 300 ms duration) overlaid with the average (black). (**D**) Population data of interictal spikes frequency versus cuprizone treatment duration. (**E**) Power spectral content during two different brain states, awake and moving (am, dotted lines) and quiet wakefulness (qw, solid lines) in control (black) and cuprizone (red). Red arrow marks the significantly amplified theta band power (θ) during quite wakefulness in cuprizone mice (Cupr_qw). (**F**) Cuprizone amplifies θ power during quiet wakefulness but not during moving. (**G**) Schematic of ECoG and LFP recordings from *Mbp*^+/+^ and *Mbp*^Shi^ mice with example trace showing interictal spikes. Higher temporal resolution of interictals in *Mbp*^Shi^ mice. (**H**) Bar plot of interictal rate in *Mbp*^Shi^ mice. Data show mean ± SEM with gray lines (D) or open circles (H) showing individual mice.

### Loss of myelin impairs fast PV^+^ BC-mediated inhibition

Increased power of sensory-driven slow oscillations and epileptiform activity is also observed when PV^+^ interneurons are optogenetically silenced ^10, 25, 37^. To investigate how myelin loss affects PV^+^ interneurons we crossed the *PV-Cre* mouse line, with Cre recombinase expression limited to *Pvalb* expressing cells, crossed with tdTomato (*Ai14*) reporter mice (*PV-Cre*; *Ai14*). The cytoplasmic fluorescence allowed quantification of PV^+^ cell bodies and their processes in the primary somatosensory cortex (**Fig. 2a, b** and Supplementary **Fig. S2a**) and immunofluorescent labelling with myelin basic protein (MBP) revealed substantial myelination of PV^+^ axons (80.15 ± 9.95% along 83 mm of PV^+^ axons analyzed, *n* = 3 slices from 2 mice, *z*-stack with a volume of 7.66 × 10^5^ μm^3^, (**Fig. 2b** and **Fig. S2a**). Electron microscopy (EM) immunogold-labeled tdTomato showed multi-lamellar compact myelin sheaths (on average, 6.33 ± 0.80 myelin lamella) with 10.8 ± 0.76 nm distance between the major dense lines and a mean *g*-ratio (axon diameter/fiber diameter) of 0.74 ± 0.01 (*n* = 6 sheaths, **Fig. 2c**). *PV-Cre; Ai14* mice fed with 0.2% cuprizone for 6 weeks showed strongly reduced MBP in S1 and PV^+^ axons were largely devoid of myelin (**Fig. 2a-b** and **Fig. S2b**) while the number of PV^+^ cells across the cortical layers remained constant (control, 326 ± 14 cells mm^−2^ vs. cuprizone, 290 ± 48 cells mm^−2^, *n* = 6 sections from *N* = 6 animals/group, Mann-Whitney test *P* = 0.1649, **Fig. 1d**). Biocytin-filled PV-Cre^+^ interneurons were re-sectioned and stained for MBP to identify the location of myelin and the axon morphology (**Fig. 2e** and **Fig. S2c, d**). Myelin was present on multiple and proximal segments of all control BCs (4/4), not on BCs from cuprizone treated mice (0/6), and the total number of axon segments (∼80 per axon, Mann-Whitney test, *P* = 0.3032, **Fig. 2f**) as well as the total path length were unaffected by cuprizone treatment (nearly ∼4 mm in each group, Mann-Whitney test, *P* = 0.9871, **Fig. 2g**, Supplementary **Fig. S2c-f**).

**Fig. 2.**
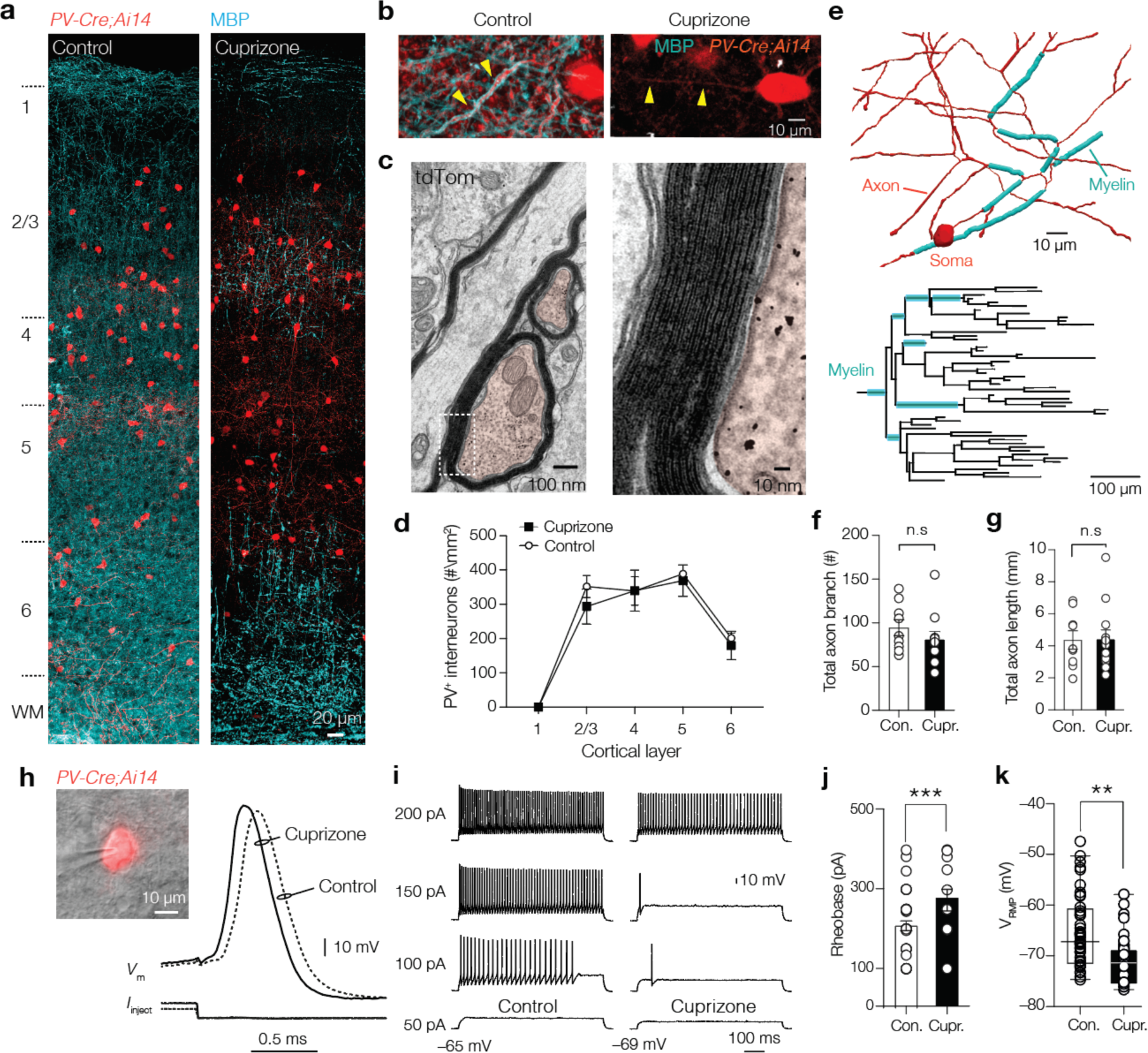
Demyelination preserves PV^+^ interneuron number and morphology but reduces excitability. (**a**) *Left*, confocal z-projected overview image of S1 in *PV-Cre*; *Ai14* animal (tdTomato^+^, red) overlaid with myelin basic protein (MBP, cyan). *Right*, overview image showing loss of MBP after 6 weeks cuprizone. (**b**) myelinated PV^+^ axons in control (*left*) and PV^+^ axons demyelination with cuprizone treatment (*Right*). (**c**) EM of transverse cut tdTomato^+^ immunogold-labelled axons (false colored red). *Right*, higher magnification of immunogold particles and ultrastructure of the PV interneuron myelin sheath. (**d**) PV^+^ interneuron number was not affected by cuprizone treatment. (**e**) *Top*, example of a high-resolution 3D reconstruction of a biocytin-labelled PV axon (red) labelled with MBP (cyan) of a control mouse, showing the first ∼6 axonal branch orders. *Bottom*, control axonogram showing axon branch order and myelinated segments (cyan). (**f-g**) Total axon branch number and length are not changed by demyelination. (**h**) *Left*, brightfield/fluorescence overlay showing patch-clamp recording from a tdTomato^+^ interneuron. *Right*, example PV^+^ interneuron APs from control (dotted line) and cuprizone treated mice (continuous lines). (**i**) Steady-state sub- and supra-threshold voltage responses during 700 ms current injections. Note the reduced firing rate near threshold. and (**j**) increased rheobase current in cuprizone. (**k**) ∼4 mV hyperpolarized resting membrane potential in demyelinated PV BCs. Data show mean ± SEM and open circles individual cells. n.s., not significant

To examine whether myelin loss changes PV^+^ BC excitability, we made whole-cell recordings in slices from *PV-Cre*; *Ai14* mice (**Fig. 2h**). Recording of steady-state firing properties by injecting increasing steps of currents injections revealed an increase in the rheobase (∼90 pA, **Fig. 2i-j**) and ∼50 Hz reduced firing frequency during low amplitude current injections (two-way ANOVA, treatment *P* = 0.0441, Šidák’s multiple comparison *post hoc* test at 200 pA; *P* = 0.0382, 250 pA; *P* = 0.0058, 300 pA; *P* = 0.0085) without a change in the maximum rates (two-way ANOVA, Šidák’s multiple comparison *post hoc* test, *P* = 0.92, data not shown) and neither the AP half-width nor amplitude (*P* = 0.7113 and *P* = 0.4358, respectively, **Table S1**). The resting membrane potential (*V*_RMP_) of control PV^+^ interneurons was on average ∼4 mV more hyperpolarized following cuprizone treatment (**Fig. 2k**, **Table S1**) without a change in the apparent input resistance (*P* = 0.5952, **Table S1**). In addition to the hyperpolarization in *V*_RMP_, demyelinated PV^+^ interneurons also had a ∼3 mV more hyperpolarized voltage threshold (Mann-Whitney test *P* = 0.0269, **Table S1**). Together, the results indicate that cuprizone treatment demyelinates PV^+^ interneuron axons without affecting anatomical properties and reduces the intrinsic interneuronal excitability.

Is myelin required for PV^+^ BC-mediated inhibition? By recording from L5 PNs several indications showed a putative impaired net inhibition. The miniature inhibitory postsynaptic currents (mIPSCs) recorded at the soma of L5 PNs were significantly reduced in peak amplitude (from ∼20 to ∼7 pA, *P* = 0.002) without a change in frequency (Supplementary **Fig. S3a–c**). Furthermore, the number of PV^+^ puncta was 40% reduced both at the somatic sites and along the primary apical dendrite and accounted to a large extent for the overall reduction in immunofluorescent signals (Supplementary **Fig. S3d-i**). Interestingly, in contrast to the loss of perisomatic PV^+^ BC puncta, putative PV^+^ chandelier cell inputs identified by co-staining PV with the AIS marker ßIV-spectrin, were preserved (∼8 puncta/AIS, Mann-Whitney test *P* = 0.96, Supplementary **Fig. S3j-k**). Furthermore, staining for Syt2, a Ca^2+^ sensor protein selective for PV^+^ presynaptic terminals ^11, 38^ similarly showed a ∼35% reduction (**Fig. S4a-b**). Syt2^+^ puncta analysis in the dysmyelinated *shiverer* mouse line also showed a reduced number of Syt2^+^ puncta at L5 PN somata and a reduced frequency of mIPSCs (*P* = 0.019, **Fig. S4c-g**), indicating that compact myelin is required for both maintaining as well as developing PV^+^ BC presynaptic terminals.

Single PV^+^ BCs typically make 5 to 15 synapses with a PN in a range of < 200 µm, forming highly reliable, fast and synchronized release sites ^5, 14, 22, 39^. We hypothesized that in addition to reduced excitability (**Fig. 2**) and PV^+^ BC release sites (Supplementary **Figs. S3** and **S4**) APs may fail to forward propagate along PV^+^ BC axons. To examine this question directly, we made paired recordings of PV^+^ BCs and L5 PNs in *PV-Cre*; *Ai14* mice, evoking APs in PV^+^ interneurons while recording unitary inhibitory post-synaptic currents (uIPSCs) in L5 PNs under conditions of physiological Ca^2+^/ Mg^2+^ (2.0/1.0 mM in *n* = 78 pairs, **Fig. 3a, b**). Concordant with optogenetic mapping of PV^+^ inputs onto L5 PNs in mouse S1 ^39^, the probability of a given PV^+^ cell being connected to a nearby PN was high (∼0.48, **Fig. 3c**). In contrast, the connection probability was significantly lower in cuprizone-treated mice (∼0.23, Chi-squared test *P* < 0.0182, **Fig. 3b, c**). In thirteen stable connected pairs, we examined unitary IPSC properties including failure rate and amplitude, as well as rise- and decay time, using automated fits of the uIPSCs (*n* > 80 trials per connection, **Fig. 3d**). Cuprizone treatment led to a significant increase in the number of failures (from 0.05 to 0.26, **Fig. 3e**) and a ∼2.5-fold reduction in the average uIPSC peak amplitude (**Fig. 3f**). Finally, to obtain an estimate of propagation speed, we determined on successful trials the latency between the AP peak and uIPSCs at 10% peak amplitude (**Fig. 3d**). Interestingly, both the mean latency was unchanged between groups (∼800 µs; Mann-Whitney test *P* > 0.999, **Fig. 3f-g**) as well as the trial-to-trial latency variability (SD in cuprizone 319 ± 65 µs, *n* = 7 pairs, SD in control, 276 ± 38 µs, *n* = 5 pairs, *P* > 0.60).

**Fig. 3.**
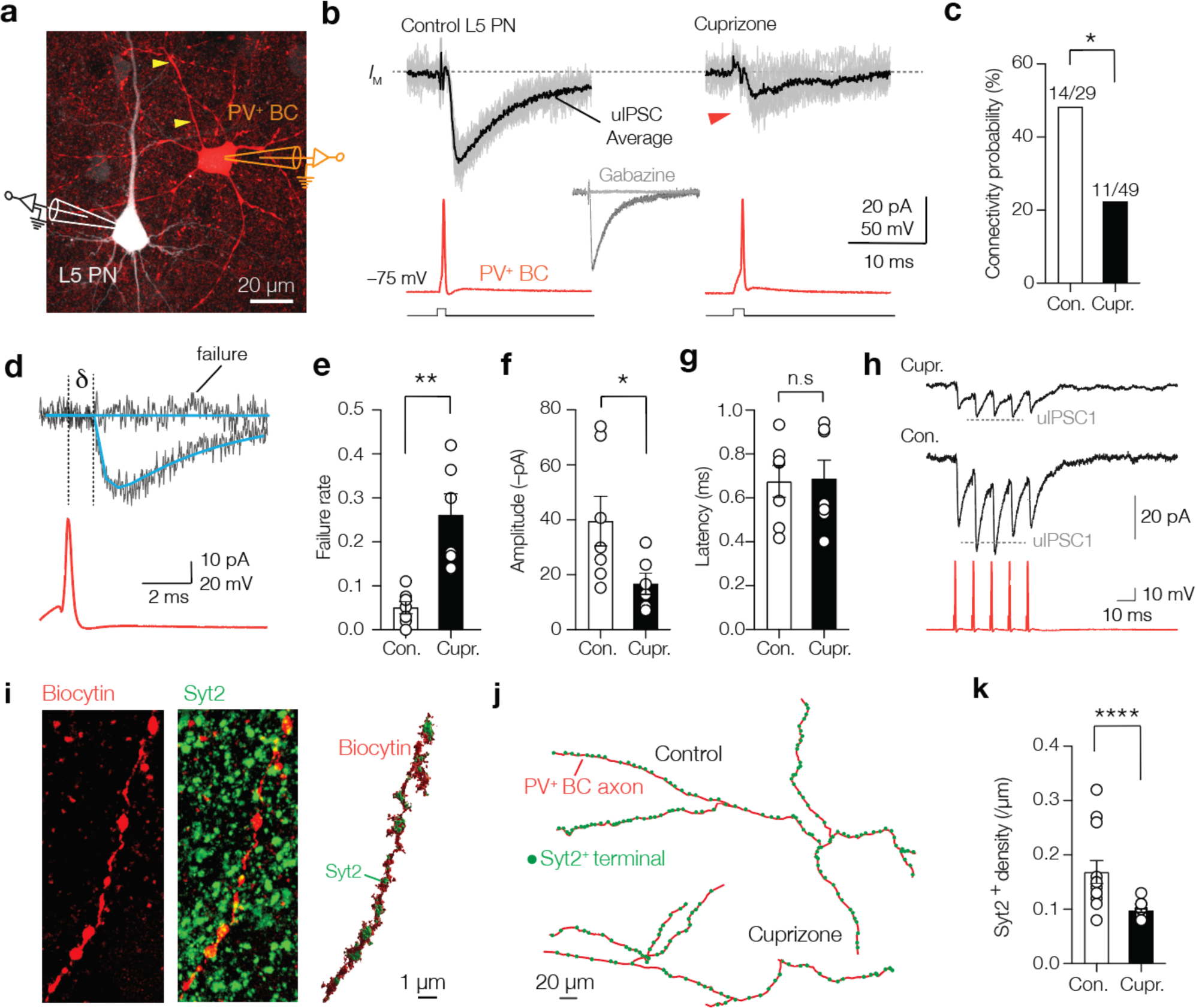
Decreased reliability, connectivity, and facilitation of PV^+^ unitary IPSCs. (**a**) Immunofluorescence image of a connected control PV^+^ BC (red) and L5 PN (white). (**b**) Example traces of ten single trial uIPSC traces (gray) overlaid with mean average (>60 trials, black). Inset, uIPSCs abolished by gabazine (GABA_A_ blocker, 4 µM). (**c**) Cuprizone-treated mice show significantly lower connection probability between PV^+^ BC and L5-PN. (**d**) Example fits (blue) of uIPSCs for rise- and decay time, amplitude, failure rate, amplitude and latency analyses (δ, AP to 10% uIPSC peak amplitude). (**e, f**) Cuprizone increased failures by 5-fold and the average peak amplitude by ∼2.5-fold. (**g**) uIPSCs latency remained unchanged. (**h**) Cuprizone impairs short-term facilitation. Dotted line indicates expected amplitude for uIPSC_2_ (scaled from uIPSC_1_). (**i**) *Left*, confocal *z*-projected image of a control PV^+^ axon (red) immunostained with Syt2 (green). *Right*, surface rendered 3D-image of the same axon. **(j**) Example sections of 3D reconstructions. (**k**) Cuprizone reduced Syt2^+^ bouton density by ∼2-fold. Data shown as mean ± SEM and open circles individual axons or pairs. n.s., not significant.

To further examine the properties of GABA release in demyelinated PV-BCs we recorded uIPSCs during a train of five APs at 100 Hz (averaging > 50 trials, **Fig. 3h**). Like the temporary facilitation in IPSCs of adult Purkinje neurons ^40^, uIPSC recordings in control PV BCs showed that paired-pulse ratios were on the second spike facilitated by 20% (uIPSC_2_/uIPSC_1_ 1.20 ± 0.060) and gradually depressed on the subsequent spikes (3 to 5). In contrast, in cuprizone-treated mice uIPSC were depressed during the second and subsequent pulses (uIPSC_2_/uIPSC_1_ 0.89 ± 0.041, **Fig. 3h**, *P* = 0.034). The uIPSC failures and impairment of temporary facilitation may reflect failure of AP propagation, changes in release probability or a lower number of active release sites (< 5, Refs. ^5, 14^) as observed in L5 neurons (**Fig. S3**). To test whether release sites along single PV^+^ BC axons are changed we performed Syt2 immunolabeling of biocytin-filled PV^+^ BCs (**Fig. 3i**). Cuprizone treatment reduced the density of Syt2^+^ puncta by 2-fold (cuprizone, ∼1 Syt2^+^ puncta per 10 µm vs. 1 Syt2^+^ puncta per 5 µm in control, Mann-Whitney test *P* < 0.0001, **Fig. 3j-k**). Thus, myelin loss reduces the presynaptic sites, increasing failures and a frequency-dependent depression limiting fast inhibition.

### PV^+^ activation rescues interictal spikes and theta oscillations, but not the loss of gamma

To examine the reduced fast PV^+^ inhibition at the network level, we used AAV1-mediated delivery of Cre-dependent channelrhodopsin-2 (ChR2) into L5 of *PV-Cre*; *Ai14* mice (**Fig. 4a, b** and Supplementary **Fig. S5a-c**). The ChR2 transduction rate was comparable between control and cuprizone mice (∼70%, **Fig. 4a**, **Fig. S5b**). In acute slices, we voltage-clamped L5 PNs and optogenetically evoked IPSC (oIPSC) with full-field blue light illumination. Consistent with S1 L5 pyramidal neurons receiving converging input from >100 PV^+^ interneurons ^39^, control oIPSCs rapidly facilitated to a peak amplitude of ∼700 pA followed by rapid synaptic depression (**Fig. 4c**). In slices from cuprizone mice, however, the oIPSC peak amplitude was ∼2-fold reduced (*P* = 0.036, **Fig. 4c**) while neither the steady-state amplitude during vesicle replenishment was not changed (**Fig. 4d**) and neither the total charge transfer reached significant difference (Control, –99.58 ± 28.5 pC vs. cuprizone, –54.3 ± 20.57 pC, *P* = 0.236, *n* = 9 control and *n* = 8 cuprizone neurons). On the other hand, miniature EPSCs recorded from PV^+^ interneurons of controlled and cuprizone-treated mice showed no changes in peak amplitude nor frequency (Supplementary **Fig. S6**), in keeping with the preservation of excitatory inputs onto L5 PNs following cuprizone-induced demyelination ^29^.

**Fig. 4.**
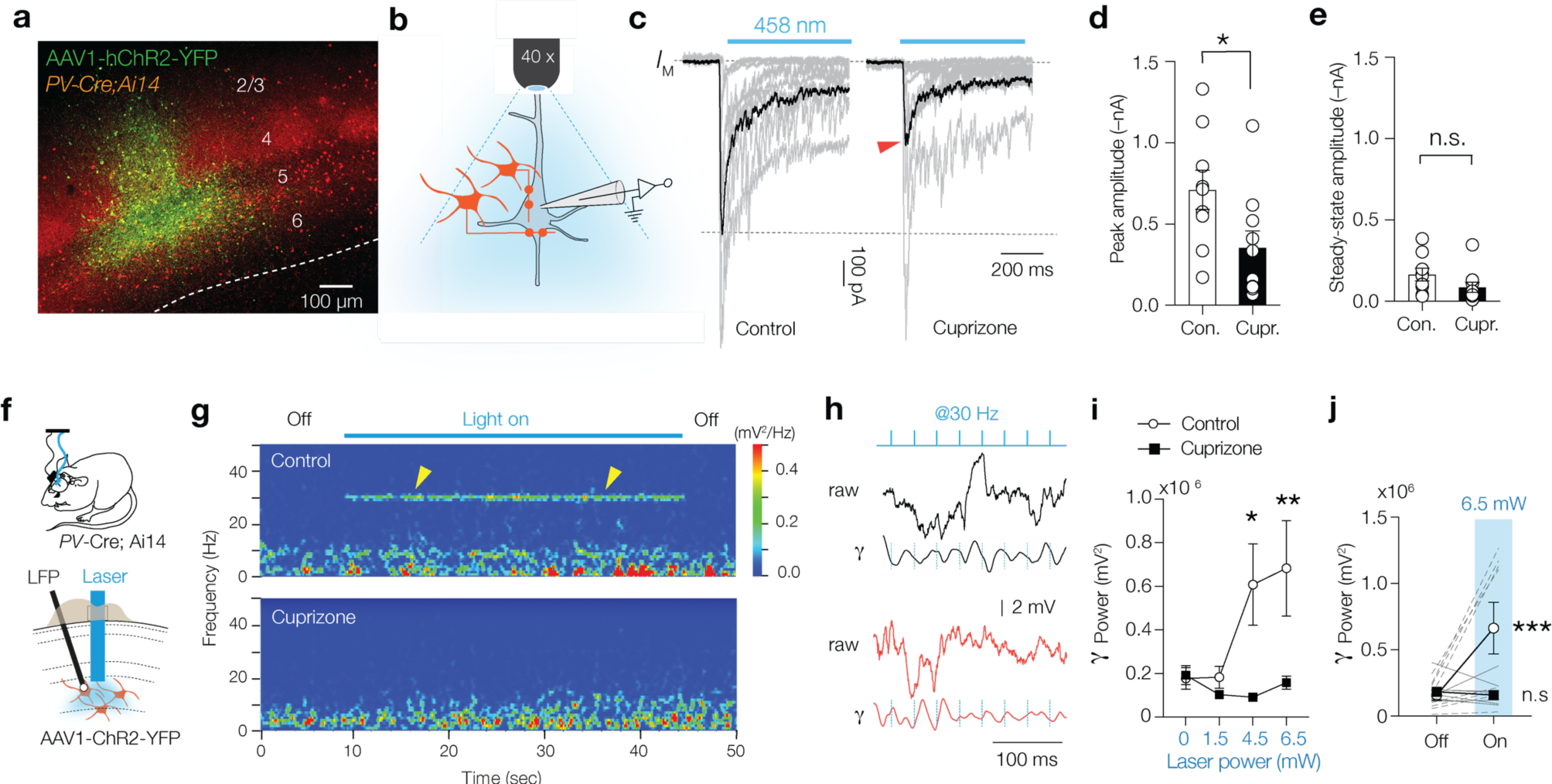
Impaired phasic PV^+^ mediated inhibition and gamma entrainment. (**a**) Immunofluorescent image of AAV1-hChR2-YFP expression (green) in L5. (**b**) schematic showing full-field blue light optogenetically-evoked postsynaptic inhibitory currents (oIPSCs) in L5 PNs. (**c, d**) Single trial oIPSCs (gray) from different experiments (1 sec duration pulses) overlaid with the average oIPSC (black), revealing a ∼2-fold reduction in oIPSCs peak amplitude (red arrow). (**e**) Steady-state oIPSCs amplitude did not reach significance. (**f**) Schematic for chronic LFP recordings and *in vivo* optogenetic stimulation in freely moving *PV-cre;Ai14* mice. (**g**) Time frequency plot showing low gamma frequency (γ) entrainment (1-ms blue light pulse at 30 Hz) in control but not in cuprizone mice. (**h**) raw LFP (*top*) and band pass filtered trace (25–40 Hz, *bottom*) from control and cuprizone during low-γ entrainment. (**i**) Population data of γ power with increasing laser power reveals impaired γ in cuprizone-treated mice. (**j**) Myelin deficient mice show lack of low-γ band during optical entrainment. Data are shown as mean ± SEM with gray lines individual cells (D, E), gray lines individual mice (J). n.s., not significant.

Impaired phasic inhibition predicts a lower power in γ rhythm ^8, 26^. Although γ power during home cage activity was not noticeably reduced (**Figs. 1e-f**) we examined the extent of γ modulation by leveraging optogenetic activation of PV^+^ interneurons with AAV1-hChR2-YFP by introducing a laser fiber into L5 and recording the LFP (**Fig. 4f**). Evoking brief pulses of blue light (1 ms, 30 Hz) showed that local circuit currents were modulated and phase-locked activity in the low γ band (between 25 and 40 Hz) of control mice (**Fig. 4g, h** and **Fig. S6**). In striking contrast, no modulation or entrainment was observed in cuprizone-treated mice (**Fig. 4i-j**, Supplementary **Fig. S6**). Could the diminished PV^+^ BC activity cause the emergence of interictal spikes during quiet wakefulness behaviors? To test the direct contribution of PV^+^ BCs in amplifying θ rhythm and interictal spikes, we activated ChR2 for 1 s duration pulses in *PV-Cre*; *Ai14* mice to generate tonic GABA release (**Fig. 5a**, **Movie S2** and Supplementary **Fig. S5c**). In cuprizone-treated mice we found that optically driving PV^+^ interneurons normalized the LFP power in the θ band to control levels, without affecting δ, β, and γ rhythms (two-way ANOVA *P* = 0.0124, light on vs. off, δ; *P* = 0.9975, θ; *P* = 0.00076, β; *P* = 0.9481, γ; *P* = 0.9998, **Fig. 5a-c**). Furthermore, activation of blue light significantly reduced the frequency of interictal epileptic discharge frequency (*P* = 0.0089, **Fig. 5d-e**). The normalization of cortical rhythms by elevating sustained PV^+^ mediated activity suggests that GABA_A_ receptors are insufficiently activated in the demyelinated cortex. Finally, to examine the role of GABA_A_ receptors agonism in dampening global interictal spikes we administered a non-sedative dose of diazepam (2 mg/kg i.p.), an allosteric modulator of post-synaptic GABA_A_ receptors, in cuprizone-treated mice (8-week treatment). The results showed that diazepam significantly suppressed the interictal epileptiform discharges in cuprizone mice (**Fig. 5f** and Supplementary **Fig. S7a, b**).

**Fig. 5.**
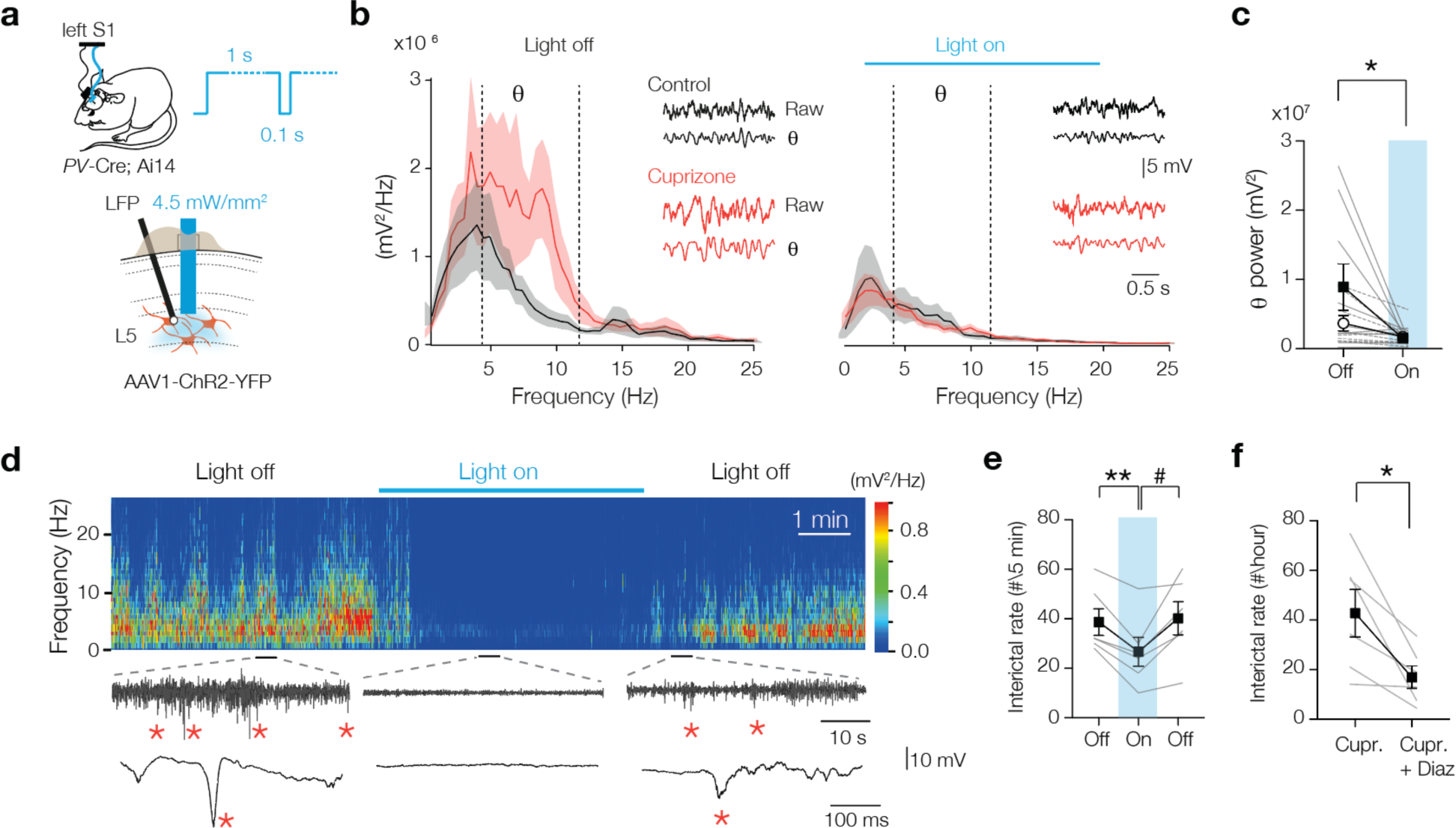
Optogenetic activation of myelin deficient PV^+^ interneurons rescues theta rhythm and interictal epileptiform discharges. **(a)** Schematic drawing for chronic LFP and optogenetic stimulation in freely moving mice. A 1 sec blue light pulse with 100 ms off periods activated PV^+^ interneurons. Blue light was switched on during high interictal activity (>10 spikes/min). (**b**) power spectral content collected from 2 sec epoch windows in control (black) and cuprizone (red) before (left) and during 3 min (right) optogenetic activation of PV^+^ interneurons. *Insets*, raw LFP signals (top) and θ content (bottom) in control (black) and cuprizone (red) condition. (**c**) Population data showing amplified θ frequency content suppressed to control levels during optogenetic activation of PV^+^ cells. (**d**) Example time frequency plot (top) and raw LFP traces (below) showing suppression of interictal spikes during light on conditions. See also **Movie S2**. (**e**) Population data of transient optogenetic suppression of the interictal activity in 8 weeks-cuprizone treated mice. (**f**) 2 mg/kg i.p injection of diazepam (diaz) in cuprizone-treated mice reduces interictals for at least 10 hours. Data show mean ± SEM and grey lines individual mice.

## Discussion

In this study we identified that the cellular microarchitecture of myelination of PV^+^ BCs contributes to fast gamma frequencies and limits the power of slow oscillations and interictal spikes during quiet wakefulness. Interictal discharges identified as spikes on the EEG are an important diagnostic criterium in epilepsy and reflect hypersynchronized burst firing of pyramidal neurons and interneurons ^32, 33, 41^. The brief episodic and generalized nature of EEG spikes we recorded in both demyelinated and dysmyelinated cortex (∼50 to 500 ms at ∼1/minute) resemble interictal spikes reported in epilepsy models ^41, 42^ and are concordant with recordings in the hippocampus of cuprizone-treated mice by Hoffmann et al. ^34^. Here we extend our insights into interictal spikes by showing spatiotemporal synchronization across cortical areas and hemispheres and their selective manifestation during the vigilance state of quiet wakefulness. During brain states of quiescence, for example when whiskers are not moving, whole-cell *in vivo* recordings in the barrel cortex reveals large-amplitude membrane potential fluctuations of PN and interneurons temporally highly synchronized at a low frequency (<10 Hz) and fast-spiking PV^+^ interneurons are dominating action potential firing ^43, 44^. These slow rhythms are internally generated and their selective amplification during quiet wakefulness is consistent with our finding of reduced intrinsic excitability of PV^+^ BCs and deficiency of fast inhibitory transmission in the demyelinated cortex (**Figs. 2, 3**). Consistent with the notion that PV^+^ disfunction suffices to amplify low frequency oscillations, in the normally myelinated cortex optogenetic inhibition of PV^+^ interneurons causes an increase of PN firing rates, elevating the power of slow oscillations and triggers epileptiform activity ^10, 25, 37^. A major limitation of the experimental toolbox available to investigate myelination is the lack of axon- or cell-type selectivity. Whether amplified delta- and abolished gamma-frequency oscillations are the consequence of PV^+^ axon demyelination, the loss of excitatory axon myelination or the combination thereof, remains to be further examined when more refined methods become available to interrogate oligodendroglial myelination of specific cell types. In the absence of such strategies, however, our *in vivo* experiments optogenetically driving PV^+^ interneurons or activating GABA_A_ receptors (**Fig. 5**) uncovers important evidence for an interneuron origin of the amplified slow oscillations and epileptic discharges. Interestingly, in contrast to dampening theta oscillations driving myelin-deficient PV^+^ interneurons at gamma-frequencies did not entrain local field oscillations. This may suggest that for gamma precisely timed spike generation of PV^+^ BCs alone is insufficient and further requires a specific circuit connectivity, GABA release dynamics or require myelination of the proximal arbors which is lost in cuprizone-treated mice. In future studies this could be examined by exploring whether remyelination reinitiates the ability of PV^+^ interneurons to produce gamma frequencies.

### Myelination of PV^+^ axons determine synapse assembly and maintenance

The prominent role of myelination of PV^+^ BCs to shape brain state-dependent rhythms is surprising in view of its sparse distribution in patches of ∼25 µm and being distributed across <10% of the total axon length ^16, 22, 45^ (**Fig. 2** and **S2**). The sparseness of interneuron myelination raised the question whether myelin speeds conduction velocity in these axon types ^16, 45, 46^. In a genetic mouse model with hypermyelinated fast-spiking interneurons the calculated conduction velocity is reduced ^24^. Here we find that the average uIPSC latency (∼800 µs) was similar between myelinated and completely demyelinated axons, and well within range of previous paired recordings between normally myelinated PV^+^ BC and PNs (700–900 µs, ^47, 48^). Assuming a typical axonal path length of ∼200 µm between the AIS and presynaptic terminals contacting a PN, combined with a ∼250 µs delay for transmitter release, the calculated conduction velocity would be 0.4 m s^−1^, consistent with optically recorded velocities (∼0.5 m s^−1^, ref. ^49^. Our paired recordings, made near physiological temperature (34 – 36 °C), may have had a limited resolution to detect temporal differences and are not excluding changes in the order of microseconds. In future studies other approaches will be required such as simultaneous somatic and axonal whole-cell recording ^50^ and/or high-resolution myelin analysis along single axon paths, which recently showed a correlation with conduction velocity ^22^. Another constraint is the lack of information on the nodes of Ranvier along demyelinated PV^+^ BC axons. With aberrant interneuron myelin development, including myelination of branch points the formation is of nodes of Ranvier is strongly disrupted^24^. Reorganization of nodal domains also occurs with the loss of myelin, affecting action potential propagation ^51, 52^ and how interneuron myelin loss changes the nodal channel distribution needs to be examined in future studies.

Converging evidence from the two distinct models (*shiverer* and cuprizone) shows that interneuron myelination critically determines PV^+^ release site number, dynamics and connection probability (**Fig. 3** and **S4**) and is concordant with the observed synapse loss along Purkinje axons of the *les* rat ^23^. The molecular mechanisms how myelination of proximal axonal segments establishes and maintains GABAergic terminals in the higher-order distal axon collaterals remains to be further investigated. The myelin sheath of PV^+^ interneurons contains high levels of non-compact 2’,3’-cyclic nucleotide 3’-phosphodiesterase (CNP) protein ^16, 53^, which is part of the inner cytoplasmic inner mesaxon ^54^. One possible mechanism may be that in the absence of inner cytoplasmic loops of oligodendroglial myelin, interneuron axons lack sufficient trophic support ^55^. In support of this idea, amyloid precursor protein, a marker of disrupted axonal transport, has been observed in early phases of cuprizone treatment and in multiple sclerosis (MS) ^56–58^. Another possibility is the pruning of GABAergic synaptic terminals by microglia ^59–61^. Microglia become increasingly activated already in sub-demyelinating stages within the first week of cuprizone treatment ^62, 63^, and in aged *Mbp*^+/–^ mice ^64^. In future studies it needs to be examined whether attenuation of microglia activation could protect against PV^+^ synapse loss and interictal epileptiform discharges.

### Implications for cognitive impairments in gray matter diseases

The identification of a cellular origin for interictal spikes may shed light to the role of PV^+^ axon myelination in cognitive impairments in MS ^65^ and possibly other neurological disorders. In models of epilepsy and patients interictal spikes have been closely linked to disruptions of the normal physiological oscillatory dynamics such as ripples required to encode and retrieve memories ^32, 42, 66, 67^. Interictal epileptic discharges are also a prominent hallmark in other cognitive diseases, including Alzheimer ^68^. Notably, reduced gray matter myelination and oligodendroglia disruption are reported in multiple epilepsy models and recently in Alzheimer ^69, 70^. Therefore, the cellular and circuit functions controlled by PV^+^ interneurons may represent a common mechanism for memory impairments in neurological disease encompassing myelin pathology. In support of this idea, neuropathological studies in MS show a specific loss of PV^+^ interneuron synapses in both cortex and hippocampus ^60, 71^. In MS patients increased connectivity and synchronization in delta and theta band rhythms during resting state or task-related behavior have been reported ^72, 73^ and low GABA levels in sensorimotor and hippocampal areas are correlated with impairments of information processing speed and memory ^74, 75^. Taken together with the present work, enhancing PV^+^ interneuron myelination, and thereby strengthening fast inhibition, may provide important new therapeutic avenues to improve cognition.

## Methods

### Animals

We crossed *PV-Cre* (B6;129P2-*Pvalb^tm1(cre)Arbr^*/J, Stock No: 008069, Jackson laboratories) with *Ai14* reporter mice (B6;129S6-*Gt(ROSA)26Sor^tm14(CAG-tdTomato)Hze^*/J, Stock No: 007908, Jackson laboratories). For other experiments we used C57BL/6 mice (Janvier Labs, Saint-Berthevin Cedex, France). Shiverer mice were obtained from Jackson (C3Fe.SWV-*Mbp^shi^*/J Stock No: 001428) and backcrossed with C57BL/6 for >10 generations. All mice were kept on a 12:12 h light-dark cycle (lights on at 07 am, lights off at 19 pm) with ad libitum food and water. For cuprizone treatment either *PV-Cre;Ai14* or C57BL/6 male or female mice, from 7 to 9 weeks of age, were fed *ad libitum* with normal chow food (control group) or were provided 0.2% (w/w) cuprizone (Bis(cyclohexanone)oxaldihydrazone, C9012, Merck) added either to grinded powder food or to freshly prepared food pellets (cuprizone group). Cuprizone-containing food was freshly prepared during every 2^nd^ or 3^rd^ day for the entire duration of the treatment (6–9 weeks). The average maximum weight loss during cuprizone feeding was ∼11% (*n* = 31). All animal experiments were done in compliance with the European Communities Council Directive 2010/63/EU effective from 1 January 2013. The experimental design and ethics were evaluated and approved by the national committee of animal experiments (CCD, application number AVD 80100 2017 2426) and the specific experimental protocols were approved and monitored under supervision of animal welfare body (IvD, protocol numbers; NIN17.21.04, NIN18.21.02, NIN18.21.05, NIN19.21.04 and, NIN20.21.02) of the Royal Netherlands Academy of Arts and Science (KNAW).

### In vitro electrophysiology

Mice were briefly anaesthetized with 3% isoflurane and decapitated or received a terminal dose of pentobarbital natrium (5 mg kg^−1^) and were transcardially perfused with ice-cold artificial CSF (aCSF) of the composition (in mM): 125 NaCl, 3 KCl, 25 glucose, 25 NaHCO_3_, 1.25 Na_2_H_2_PO_4_, 1 CaCl_2_, 6 MgCl_2_, 1 kynurenic acid, saturated with 95% O_2_ and 5% CO_2_, pH 7.4. After decapitation, the brain was quickly removed from the skull and parasagittal sections (300 or 400µm) containing the S1 cut in ice-cold aCSF (as above) using a vibratome (1200S, Leica Microsystems). After a recovery period for 30 min at 35 °C brain slices were stored at room temperature. For patch-clamp recordings, slices were transferred to an upright microscope (BX51WI, Olympus Nederland) equipped with oblique illumination optics (WI-OBCD; numerical aperture, 0.8). The microscope bath was perfused with oxygenated (95% O_2_, 5% CO_2_) aCSF consisting of the following (in mM): 125 NaCl, 3 KCl, 25 D-glucose, 25 NaHCO_3_, 1.25 Na_2_H_2_PO_4_, 2 CaCl_2_, and 1 MgCl_2_. L5 pyramidal neurons were identified by their typical large triangular shape in the infragranular layers and in slices from *PV-Cre;Ai14* mice the PV^+^ interneurons expressing tdTomato were identified using X-Cite series 120Q (Excelitas) with a bandpass filter (excitation maximum 554 nm, emission maximum 581 nm). Somatic whole-cell current-clamp recordings were made with a bridge current clamp amplifier (BVC-700A, Dagan Corporation, US) using patch pipettes (4–6 MΩ) filled with a solution containing (in mM): 130 K-gluconate, 10 KCl, 4 Mg-ATP, 0.3 Na_2_-GTP, 10 HEPES, and 10 Na_2_-phosphocreatine, pH 7.4, adjusted with KOH, 280 mOsmol/kg, to which 10 mg mL^−1^ biocytin was added. Voltage was analog low-pass filtered at 10 kHz (Bessel) and digitally sampled at 50–100 kHz using an analog-to-digital converter (ITC-18, HEKA Electronic) and data acquisition software Axograph X (v.1.7.2, Axograph Scientific). The access resistance was typically < 20 MΩ and fully compensated for bridge balance and pipette capacitance. All reported membrane potentials were corrected for experimentally determined junction potential of –14 mV. Analysis for the electrophysiological properties includes PV interneuron recordings from cells in normal ACSF and in the presence of CNQX and d-AP5 with high chloride intracellular solution (see below).

### mIPSC and mEPC recordings

Whole-cell voltage-clamp recordings were made with an Axopatch 200B amplifier (Molecular Devices). Patch pipettes with a tip resistance of 3–5 MΩ were pulled from thins wall borosilicate glass. During recording, a holding potential of –74 mV was used. Both the slow- and fast pipette capacitance compensation were applied, and series-resistance compensated to ∼80-90%. Patch pipettes were filled with high chloride solution containing (in mM): 70 K-gluconate, 70 KCl, 0.5 EGTA, 10 HEPES, 4 MgATP, 4 K-phosphocreatine, 0.4 GTP, pH 7.3 adjusted with KOH, 285 mOsmol kg^−1^ and IPSCs isolated by the presence of the glutamate receptor blockers 6-cyano-7-nitroquinoxaline-2,3-dione (CNQX, 20 µM), d-2-Amino-5-phosphonovaleric acid (d-AP5, 50µM) and the sodium (Na^+^) channel blocker tetrodotoxin (TTX, 1 µM Tocris). Individual traces (5 sec duration) were filtered with a high-pass filter of 0.2 Hz and decimated in Axograph software. Chart recordings of mIPSCs were analyzed with a representative 30 ms IPSC template, using the automatic event detection tool of Axograph. Detected events were aligned and averaged for further analysis of inter-event intervals (frequency) and peak amplitude. For mEPSC recordings from PV^+^ interneurons we filled patch pipettes with a solution containing (in mM): 130 K-gluconate, 10 KCl, 4 Mg-ATP, 0.3 Na_2_-GTP, 10 HEPES, and 10 Na_2_-phosphocreatine, pH 7.4, adjusted with KOH, 280 mOsmol/kg and both gabazine (4 µM) and TTX (1 μM) were added to the bath solution. The mEPSCs were analyzed using events detection tool in Axograph. The recorded signals were bandpass filter (0.1 Hz to 1 kHz) and recordings analyzed with a representative 30 ms EPSC template, after which selected EPSCs aligned and averaged for further analysis of inter-event intervals (frequency) and peak amplitude.

### uIPSC recording and analysis

PV^+^ interneurons (identified in *PV-Cre;Ai14* mice) were targeted for whole-cell current-clamp recording within a radius of 50 µm from the edge of the L5 soma recorded in voltage-clamp configuration. APs in PV^+^ interneurons were evoked with a brief current injection (1–3 ms duration) and uIPSCs recorded in the L5 PN from a holding potential of –74 mV. Only responses with 2 × S.D. of baseline noise were considered being connected. Both fast and slow capacitances were fully compensated, series-resistance compensation was applied to ∼80-90% and the current and voltage traces acquired at 50 kHz. For stable recordings with > 50 uIPSCs the episodes were temporally aligned to the AP and the uIPSCs were fit with a multiexponential function in Igor Pro. The curve fitting detected the baseline, uIPSC onset, rise time, peak amplitude and decay time and was manually monitored. Fits were either accepted or rejected (e.g. when artefacts were present) and the number uIPSC failures were noted for each recording.

### In vitro optogenetics

50 nL of AAV1 particles (titer 1 × 10^12^ cfu mL^−1^) produced from pAAV-EF1a-double-floxed-hChR2(H134)-EYFP-WPRE-HGHpA (Addgene.org #20298) was injected into L5 of S1 (co-ordinates from bregma; AP-0.15 mm ML-0.30 mm and DL-0.75 mm) of 6–9 weeks old *PV-Cre; Ai14* mice. About 7 days after the injection, a subset of mice was placed on 0.2% cuprizone diet for 8 to 9 weeks. PV^+^ interneurons expressing hChR2 were identified using td-tom and YFP co-expression. Whole-cell voltage-clamp recordings were made from L5 PNs and optically induced inhibitory postsynaptic currents (oIPSCs) were evoked with a X-cite 120Q, fluorescent lamp using filter BA460-510 (Olympus) in the presence of CNQX (50 µM) and dAP5 (20 µM) in the bath solution. The oIPSCs were evoked by illumination of large field with 5 light pulses of each 1 ms and 100 ms apart. Peak amplitude and area under curve (charge) of oIPSC was quantified using Axograph. Only the first pulse was used for the quantification.

### In-vivo electrophysiology and automated event detection

Chronic ECoG and LFP recordings were performed using in-house made electrodes of platinum-iridium wire (101R-5T, 90% Pt, 10% Ir, complete diameter of 200 µm with 127 µm metal diameter, Science Products). The perfluoroalkoxy alkanes (PFA) coated wire platinum-iridium wire was only exposed at the tip to record the local field potential (LFP). For placement of the recording electrode, animals were anesthetized with isoflurane (3%, flow rate 0.8 L/min with maintenance 1.5–1.8%, flow rate 0.6 L/min). A 1 cm midline sagittal incision was made starting above the interaural line and extending along the neck to create a pocket for subcutaneous placement of the transmitter along the dorsal flank of the animal. The recording electrodes in each hemi-sphere (stereotaxic coordinates relative to bregma: S1; –0.15 mm anterior and ± 0.30 mm lateral; for LFP; ventral 0.75 mm, V1; 0.40 mm anterior and ± 0.30 mm lateral; for LFP; ventral 0.75 mm) and ground electrode (6 mm posterior and 1 mm lateral) were implanted sub-durally through small holes drilled in the skull, held in place with stainless steel screws (A2-70, Jeveka) and subsequently sealed with dental cement. Mice were provided with Metachem analgesic (0.1 mg per kg) after surgery and allowed to recover for 4–7 days before recordings. To obtain multiple hours recordings of ECoG-LFP at multiple weeks, mice remained in their home cage during an overnight recording session. ECoG–LFP data were collected using a ME2100-system (Multi channel Systems); ECoG-LFP data were acquired at a sampling rate of 2 kHz using the multi-channel experimenter software (Multi channel systems). An additional 0.1–200 Hz digital band-pass filter was applied before data analysis. Large noise signals, due to excessive locomotion or grooming, were manually removed from the data. The ECoG and LFP recordings were processed offline with the Neuroarchiver tool (Open Source Instruments, http://www.opensourceinstruments.com/Electronics/A3018/Seizure_Detection.html). To detect interictal spikes an event detection library was built as described previously ^31^. During the initial learning phase of the library the observer, if needed, overruled the identity of each new event by the algorithm, until automated detection reached a false positive rate < 1%. Subsequently, the ECoG-LFP data were detected by using a single library across all ECoG-LFP recordings. For determining the interictal rate, only S1 LFP signals were used for quantification.

### In-vivo optogenetics with simultaneous ECoG-LFP recordings

50 nl of AAV1 particles (titer 1 × 10^12^ cfu ml^−1^) produced from pAAV-EF1a-double-floxed-hChR2(H134)-EYFP-WPRE-HGHpA (Addgene #20298) was injected unilaterally into the L5 of S1 (coordinates from bregma; AP-0.15 mm ML-0.30 mm and DL-0.75 mm) of 6–9 weeks old *PV-Cre; Ai14* mice. ECoG-LFP electrode (stereotaxic coordinates relative to bregma: –0.15 mm anterior and ± 0.30 mm lateral; for LFP; ventral 0.75 mm) and ground electrode (6 mm posterior and 1 mm lateral) were implanted through small holes drilled in the skull, held in place with stainless steel screws (A2-70, Jeveka). Through the drilled hole, a polished multimode optical fiber (FP200URT, Thorlabs) held in ceramic ferrule (CFLC230-10, Thorlabs) was driven into the layer 5 and ∼50 µm above virus injection site. Once optical fiber and electrode were correctly placed, the drilled hole subsequently sealed with dental cement. A blue fiber-coupled laser (473 nm, DPSS Laser T3, Shanghai Laser & Optics Co.) was used to activate the ChR2. Cyclops LED Driver (Open ephys) together with customized program was used to design the on and off state of the laser. The driving signal from LED driver was also recorded at one of the empty channels in multi-channel systems. This signal was used to estimate the blue light on or off condition. For gamma entrainment in S1, 40 pulses of blue light were flashed with 1 ms on and 28 ms off pulse.

To inhibit interictal spikes, 300 pulses of blue light were flashed with 1 sec on and 100 ms off by manual activation of light pulses when periods of high interictal spikes were observed (> 40 interictals/min). Aged-matched control mice were stimulated during the resting phase of the EEG, which was estimated using online EMG signal and video observation. For interictal counts, 5 min LFP signals were used from before light stimulation, during, and post light stimulation. Interictal were detected using event detection library. For analysis of the cortical rhythms, epochs were extracted using 2 second window at the start and after 180 pulses of blue light. Epoch containing interictal were not included in the analysis.

For pharmacology experiment, continuous LFP recordings of > 10-12 hours duration from the circadian quiet phase (from 19:00 to 09:00) of 6 cuprizone mice (7 weeks treatment) and 3 control mice were used for the analysis. To activate GABA_A_ receptors in cuprizone-treated mice, we used diazepam (Centrafarm Nederland B.V) prepared in a 10% solution of (2-Hydroxypropyl)-β-cyclodextrin (Sigma-Aldrich). A non-sedative dose of 2 mg kg^−1^ diazepam was injected intraperitoneally, and data was acquired for a period of 10 hours, starting 15 min after injection of drug in control and cuprizone mice. The automated event detection library (**Fig. S1**) was used to determine the event frequency before and after diazepam injection.

### In-vivo power spectrum analysis

Power spectral density (PSD) analysis was done using multi-taper PSD toolbox from Igor Pro 8.0. The absence of high voltage activity in the EMG electrode was classified as quiet wakefulness (**Fig. S1** and **Movie S1**). For PSD analysis during interictal activity, a 2 sec window was used to extract LFP signal epochs. Epochs from control animals were selected comparing the EMG activity with cuprizone EMG activity. The interictal activity itself was excluded from the analysis. Selected LFP epochs were band pass filtered between different frequency bands; delta, δ, (0.5-3 Hz), theta, θ, (4-12 Hz), beta, β, (12.5-25 Hz) and gamma, γ, (30-80 Hz). Multi-taper PSD function (Igor Pro 8.0) was applied to the filtered the data to plot the power distribution within each frequency band. Areas under the curve was measured for each frequency band to compare power density between the control and cuprizone groups.

### Immunohistochemistry

L5 PNs were filled with 10 mg ml^−1^ biocytin during whole-cell patch clamp recording for at least 30 minutes. Slices were fixed for 30 min with 4% paraformaldehyde (PFA) and stored in 0.1 M phosphate buffered saline (PBS; pH 7.4) at 4 °C. Fixed 400 μm slices were embedded in 20% gelatin (Sigma-Aldrich) and then sectioned with a Vibratome (VT1000 S, Leica Microsystems) at 80 μm. Sections were pre-incubated with blocking 0.1M PBS containing 5% normal goat serum (NGS), 5% bovine-serum albumin (BSA; Sigma-Aldrich) and 0.3% Triton-X (Sigma) during 2 hours at 4 °C to make the membrane permeable. For biocytin-labelled cells, streptavidin biotin-binding protein (Streptavidin Alexa 488, 1:500, Invitrogen) was diluted in 5% BSA with 5% NGS and 0.3% Triton-X overnight at 4 °C. Sections including biocytin-filled cells were incubated again overnight at 4 °C with primary antibody rabbit anti-ßIV-spectrin (1:200; gift from M.N. Rasband, Baylor College of Medicine), mouse anti-myelin basic protein (MBP) (1:250; Covance), mouse anti-PV (1:1000; Swant) rabbit anti-syt2 (1:500, Synaptic Systems) in PBS blocking solution containing 5% BSA with 5% NGS and 0.3% Triton-X. Secondary antibody were used to visualize the immunoreactions: Alexa 488-conjugated goat anti-rabbit (1:500; Invitrogen), Alexa 488 goat anti-mouse (1: 500; Sanbio), Alexa 488 goat anti-guinea pig, Alexa 555 goat anti-mouse (1:500; Invitrogen), Alexa 555 goat anti-rabbit (1:500; Invitrogen), Alexa 633 goat anti-guinea pig (1:500; Invitrogen), Alexa 633 goat anti-mouse (1:500; Invitrogen) and Alexa 633 goat anti-rabbit (1:500; Invitrogen). Finally, sections were mounted on glass slides and cover slipped with Vectashield H1000 fluorescent mounting medium (Vector Laboratories, Peterborough, UK) and sealed.

### Confocal imaging

A confocal laser-scanning microscope SP8 X (DM6000 CFS; acquisition software, Leica Application Suite AF v3.2.1.9702) with a 63× oil-immersion objective (1.3 NA) and with 1× digital zoom was used to collect images of the labelled L5 neurons and the above-mentioned proteins. Alexa fluorescence was imaged using corresponding excitation wavelengths at 15 units of intensity and a *z*-step of 0.3 μm. Image analysis was performed with Fiji (ImageJ) graphic software (v.2.0.0-rc-65/1.5w, National Institutes of Health). Putative PV^+^ puncta counting or Syt2 was manually done by trained personal blinded to the identity of the experimental groups.

### Synaptic puncta counting and image analysis

The intensity of PV^+^ or Syt2 immunostaining was measured with a *z*-axis profile, calculating the mean RGB value for each *z*-plane. In quantifying the axosomatic projections, the soma is defined by cutting off the apical dendrite at ∼4 μm from an imaginary rounding of the soma. The boutons were selected by hand indicated either by colocalization of the pyramidal cell and PV/Syt2 or direct contact of the two. The boutons were characterized as round spots with a minimal radius of 0.5 μm ranging to almost 2 μm. All image analysis was done in FIJI (ImageJ) graphic software (v2.0, National Institutes of Health).

### PV^+^ axon reconstruction and quantification

For immunolabeling of biocytin-filled PV^+^ interneuron, 400 μm electrophysiology slices were incubated overnight at 4 °C in PFA. Slices were rinsed with PBS followed by staining using streptavidin 488 (1:300, Jackson) diluted in PBS containing 0.4% Triton-X and 2% normal horse serum (NHS; Gibco) overnight at 4 °C. Confocal images of 400 μm thick slices were taken (see Methods, Confocal Imaging) and immediately after, thoroughly rinsed with 0.1M PB and 30% sucrose at 4 °C overnight. Next, slices were sectioned into 40-μm thick and preserved in 0.1 M PB before staining. Sections were pre-incubated in PBS blocking buffer containing 0.5% Triton-X and 10% NHS during one hour at room temperature. Sections were stained with primary mouse anti-MBP (1:300, Santa Cruz), rabbit anti-syt2 in 0.4% Triton-X and 2% NHS with PBS solution for 72 h. Alexa 488-conjugated secondary antibodies (1:300, Invitrogen) were added in PBS containing 0.4% Triton-X and 2% NHS, posterior to washing steps with PBS. Then, sections were mounted on slides and cover slipped with Vectashield H1000 fluorescent mounting medium, sealed and imaged. Biocytin-labelled PV^+^ neurons were imaged using upright Zeiss LSM 700 microscope (Carl Zeiss) with 10× and 63× oil-immersion objectives (0.45 NA and 1.4 NA, respectively) and 1× digital zoom with step size of 0.5 µm. Alexa 488 and Alexa 647 were imaged using 488 and 639 excitation wavelengths, respectively. The 10× image was taken to determine the exact location of biocytin-filled cells. Subsequently, axonal images were taken at 63× magnification. Axons were analyzed as described previously ^45^ and identified by their thin diameter, smoothness, obtuse branching processes and occasionally by the presence of the axon bleb. Images were opened in Neurolucida 360 software (v2018.02, MBF Bioscience) for reconstruction using the interactive user-guided trace with the Directional Kernels method. Axon and myelinated segments were analyzed using Neurolucida Explorer (MBF Bioscience). Axonal segments were accepted as myelinated when at least one MBP-positive segment co-localized with streptavidin across the internode length.

### Statistics

For comparisons of two independent groups, we used two-tailed Mann-Whitney U tests. For multiple groups comparisons, data were assessed for normality and either an ordinary two-way analysis of variance (ANOVA) or two-way ANOVA with repeated measures followed by Šidák’s multiple comparisons test was applied using Prism 8 (Version 8.3.0, GraphPad Software). The level of significance was set to 0.05 for rejecting the null hypothesis. An overview of the results from all statistical analyses is presented in **Table S2**.

## Data availability

All raw data will be made available upon request to the corresponding author.

## Acknowledgements

The authors are indebted to Prof. Dr. Stefan Hallermann (University of Leipzig) for providing the uIPSC analysis script. We thank Ms. Anouk Meuwissen, Catherine Jenkins, Denise de Ronde and Dr. Koen Kole (NIN–KNAW) with their support in part of the recordings and optogenetic experiments. Sharon I. De Vries performed the electron microscopy. Dr. Corette Wierenga (UU) and Dr. David Vandael provided highly valuable comments on earlier versions of the manuscript and experimental work. This work was in part funded by The National Multiple Sclerosis Society RG-1602-07777 (M.K.), The Netherlands Research Council NWO Vici 865.17.003 (M.K.), The Netherlands Research Council NWO 013.18.002 (S.A.K.), European Research Area Network ERA-NET NEURON JTC2018-024 (S.A.K.)

## Author contributions

Conceptualization: M.K., Visualization: M.K., M.D., Experimental design: M.K., M.D., Supervision: M.D., S.K., M.K., Funding acquisition: M.K., S.K., Methodology: M.D., M.K., S.K., Software: M.D., Investigation: M.D., M.G., K.H., D.W., M.H., M.K., Formal analysis, M.D., M.G., M.K., Writing – original draft, M.K., Writing – reviewing & editing final draft, M.D., M.K., S.K., Data curation, M.D., M.K.

## Materials & correspondence

Requests should be directed to Prof. Dr. Maarten H.P. Kole

## Supplementary information

**Fig. S1.**
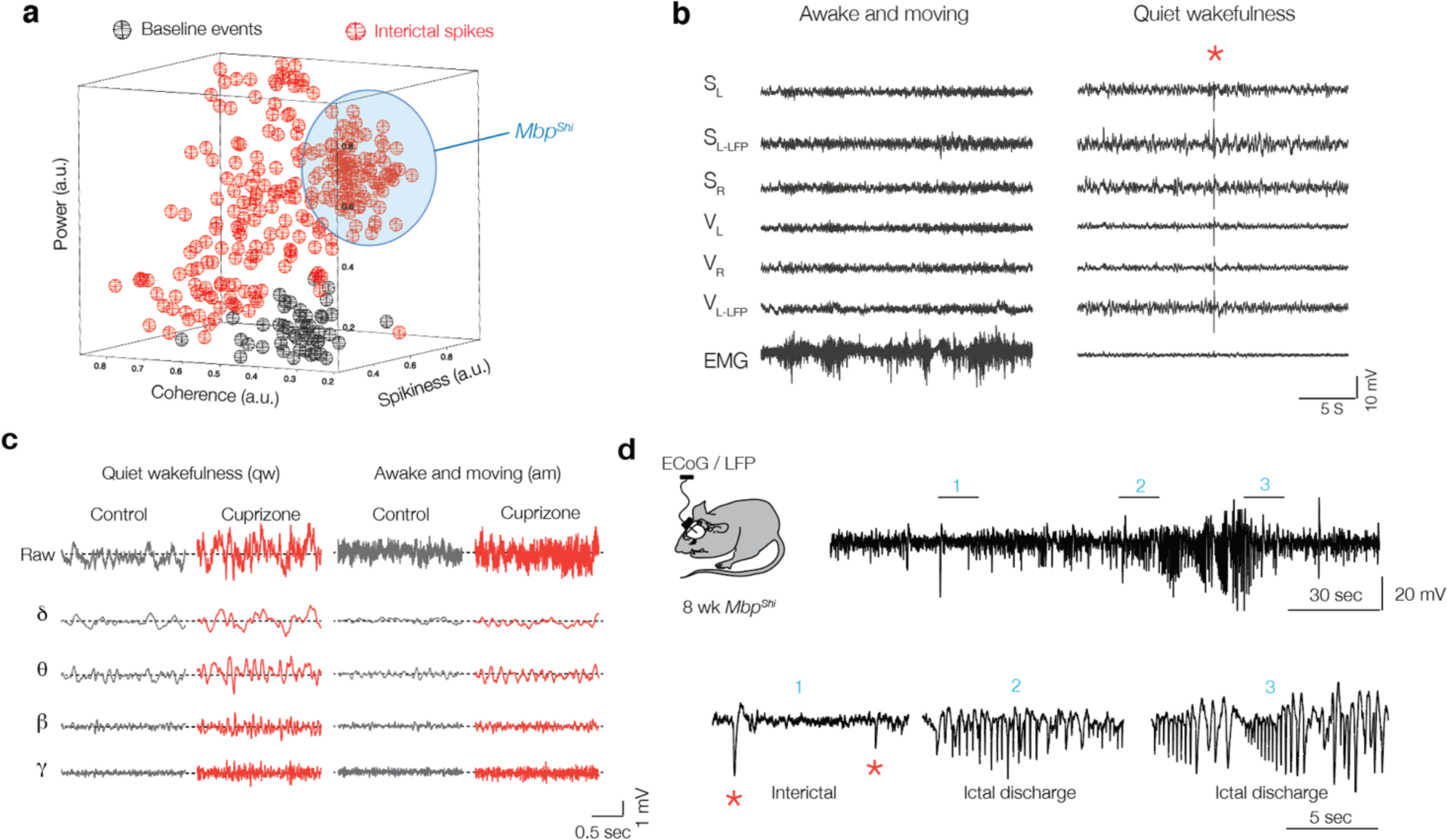
State-dependent interictal activity in cuprizone mice and automated interictal event detection library. (**a**) Automated event detection library used for interictal classification. Three-dimensional projection metric space, showing coherence, spikiness and signal power (a.u. = arbitrary units), with colors corresponding to interictal (red) and baseline/normal (black) events. The event library was constructed by an operator who classified events as “normal” or interictal events. The blue area represents the population of interictal events from *Mbp*^Shi^ mice. (**b**) Example of a 30 s recording from multiple electrodes from 6 weeks-cuprizone treated mouse. *Left*, traces during the awake state, note the high voltage EMG activity. *Right*, same mouse during quiet wakefulness with low EMG activity. Interictal discharge indicated with red asterisk (*). (**c**) Example traces showing raw LFP signals from S1 (top) and bandpass filtered traces (bottom) at different brain states in control (black) and cuprizone (red) for delta (δ, 0.5-3.5 Hz), theta (θ, 4-12 Hz), beta (β, 12.5-25 Hz) and gamma (γ, 30-80 Hz). (**d**) Schematic drawing showing ECoG and LFP recordings from S1 of *Mbp*^+/+^ and *Mbp*^Shi^ mice. Example LFP trace showing pre-ictal and ictal discharge from 8 weeks old *Mbp*^Shi^ mouse (top). Bottom, higher temporal resolution from the top LFP trace showing pre-ictal discharge with interictals (1) and ictal discharge (2 and 3).

**Fig. S2.**
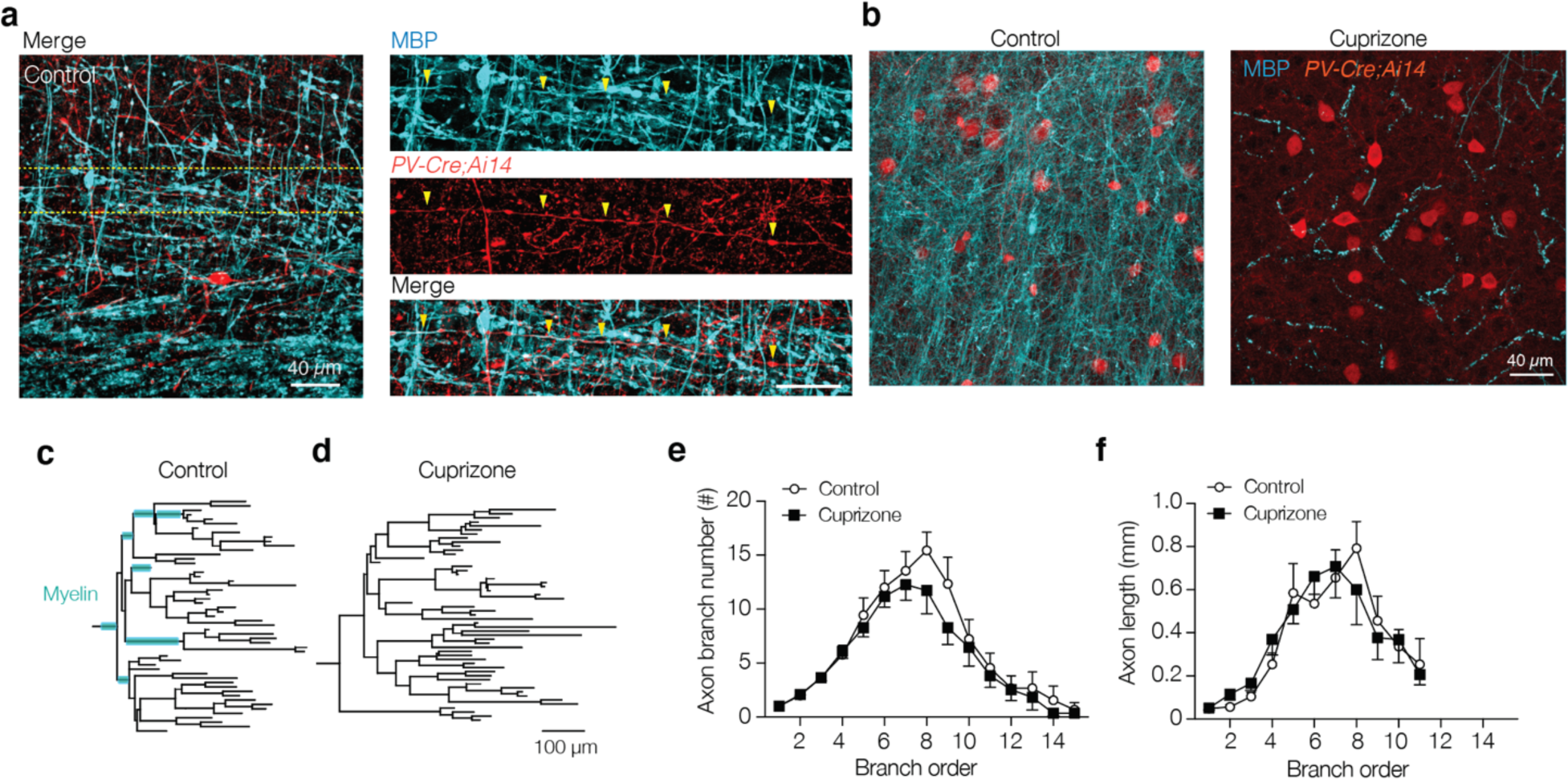
Cuprizone-induced demyelination preserves PV^+^ axon length and complexity. (**a**) *Left*, confocal image of control staining in L5 of S1 showing a PV^+^ interneuron. *Right*, higher magnification of the image in *right*, illustrating the trajectory of a myelinated PV^+^ interneuron axon (yellow arrows). Note that also PV^+^ axon swellings are frequently myelinated. Scale bar, 10 µm. (**b**) A confocal *z*-stack image examples of the L5 region in control (left) and cuprizone (right) treated mice. (**c**) Axonograms of a control (left) and cuprizone axon (**d**, right). Myelinated segments indicated with cyan. (**e-f**) Number/length per branch order. Axon branch number and segment lengths are not affected with cuprizone-induced demyelination (*P* = 0.8028 and *P* = 0.6236, respectively). Data show mean ± SEM.

**Fig. S3.**
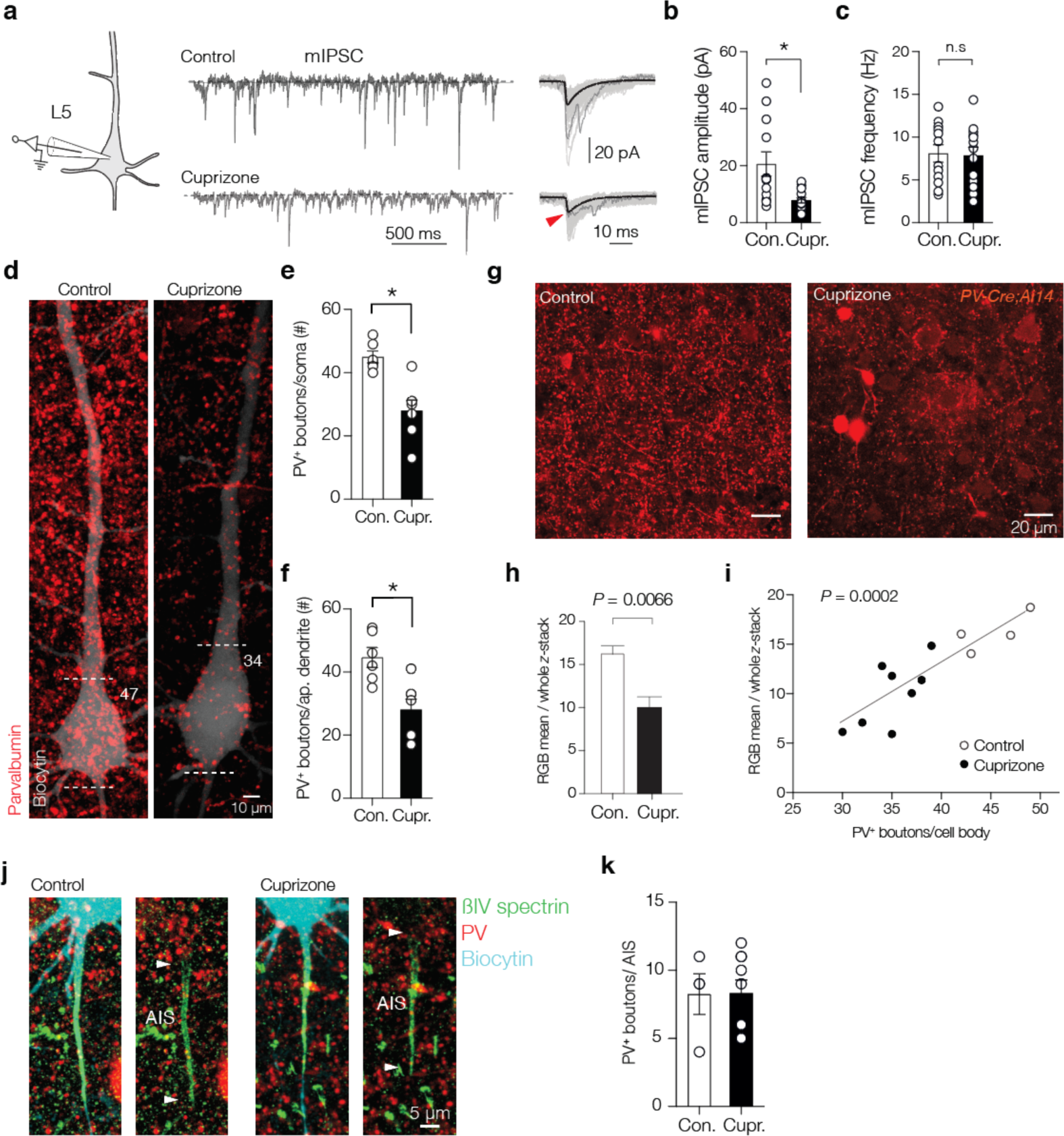
Cuprizone decreases miniature IPSCs and somatodendritic PV puncta numbers. (**a**) *Left*, Example traces of mIPSCs at the soma of L5 pyramidal neurons in the presence of CNQX, d-AP5 and TTX in control (top) and demyelinated conditions (bottom). (**b**) Population data showing a ∼3-fold mIPSC peak amplitude reduction. (**c**) Miniature frequency was unaffected. (**d**) Maximum *z*-projection of a biocytin filled L5 pyramidal neurons (white) overlaid with PV immunofluorescence (red). Dotted lines indicate the soma borders with number indicating PV^+^ puncta. (**e**) Population data showing significant loss in the number of PV^+^ puncta at the L5 soma. (**f**) Population analysis of PV^+^ puncta at primary proximal apical dendrite (< 200 µm) shows a reduced puncta number. (**g, h**) Example confocal *z*-stack images of L5 in control (left) and cuprizone treated mice (right) reveals a global significant reduction in PV immunofluorescence intensity. (**i**) Regression plot reveals the mean PV immunofluorescence intensity correlates with the number of PV^+^ boutons on large NeuN^+^ pyramidal neurons cell bodies (*r*^2^ = 0.755). NeuN^+^ immunofluorescent signals are not shown. The soma diameters of pyramidal neurons were unchanged (cuprizone 18.23 ± 0.99 µm, *n* = 6 vs. 18.37 ± 0.68 µm, Mann-Whitney U test *P* = 0.954, *n* = 9). (**j**) A confocal *z*-projection of a biocytin filled (cyan) L5 pyramidal neuron overlaid with βIV spectrin (green). (**k**) putative chandelier PV^+^ boutons were preserved in cuprizone-treated mice. Data are shown as mean ± SEM with open circles individual neurons. n.s., not significant.

**Fig. S4.**
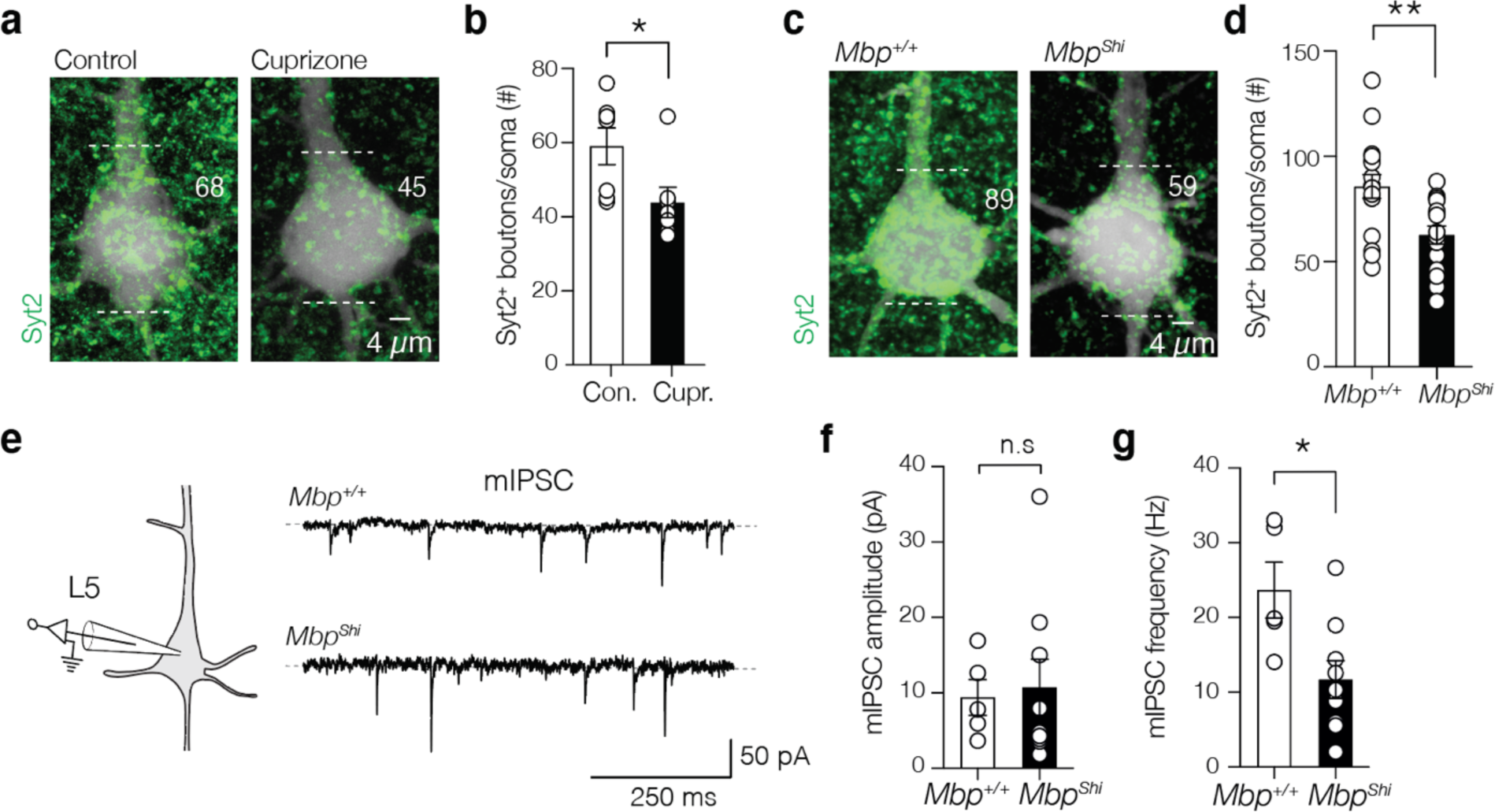
Demyelination and dysmyelination reduces miniature IPSCs and perisomatic Syt2^+^ puncta. (**a**) Maximum *z*-projection of biocytin-filled L5 PN (white) overlaid with Syt2^+^ immunofluorescence (green). Numbers indicate the Syt2^+^ puncta. (**b**) Population analysis of Syt2^+^ puncta reveal a significant loss in cuprizone. (**c**) Maximum *z*-projection of a biocytin filled L5 soma (white) overlaid with Syt2^+^ immunofluorescence (green) from *Mbp*^+/+^ and *Mbp*^Shi^ mice. (**d**) Population analysis shows a significant loss of Syt2^+^ puncta in the *Mbp*^Shi^ mice. (**e**) Example traces of mIPSCs of L5 PNs in the presence of CNQX, d-AP5 and TTX in *Mbp*^+/+^ (top) and *Mbp*^Shi^ mice (bottom). (**f, g**) mIPSCs peak amplitude was unaffected but frequency is reduced in *Mbp*^Shi^ (**\****P* = 0.019). Data are shown as mean ± SEM with open circles individual neurons. n.s., not significant.

**Fig. S5.**
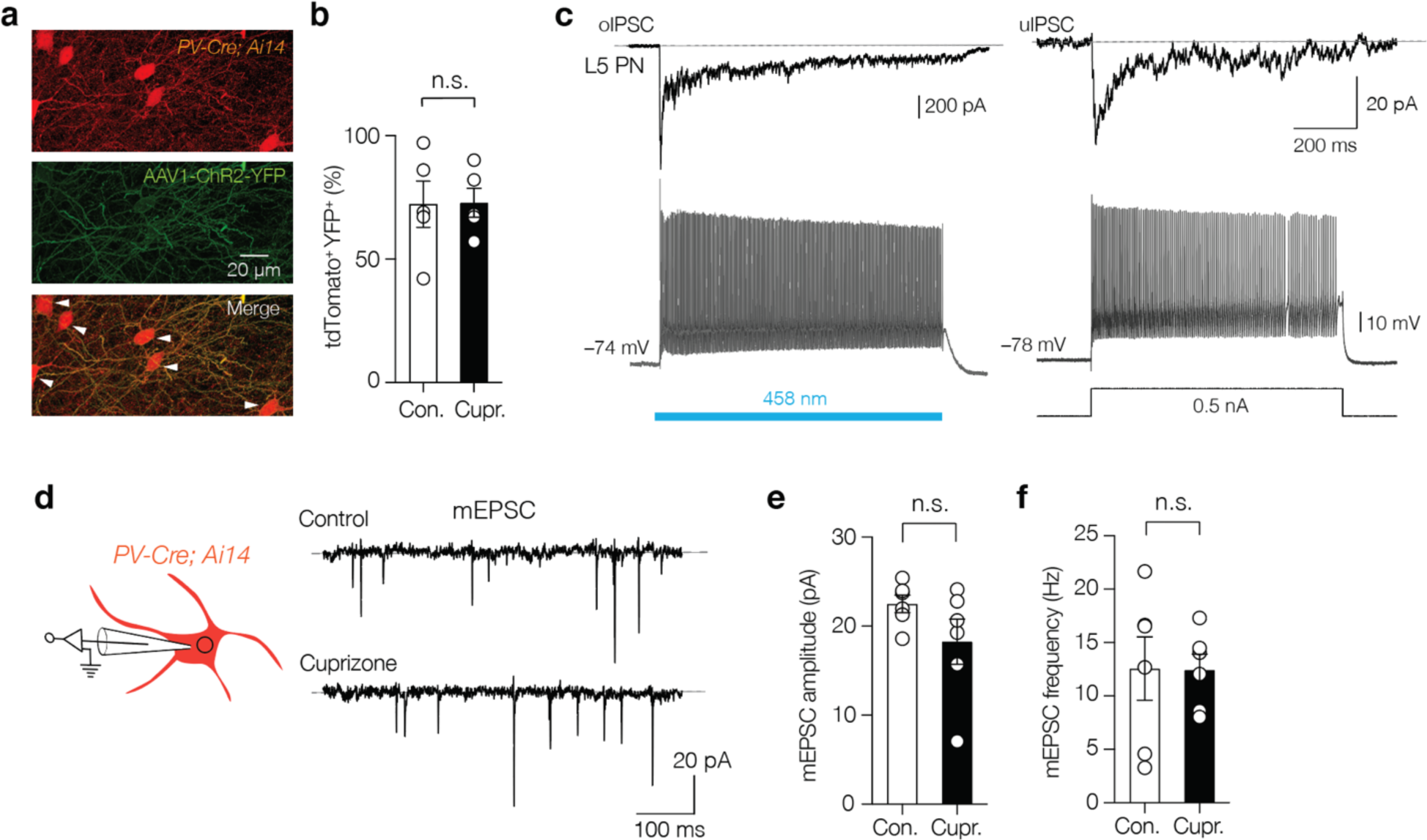
Optogenetically evoked PV^+^ IPSC resemble unitary IPSCs and excitatory drive of PV BCs remains unaffected. (**a**) confocal image of separate fluorescent channels showing td-Tomato^+^ cell bodies and neurites (red), the localization of AAV1-hChR2-YFP (green) and the merge image. The majority of tdTomato^+^ cells were YFP^+^ (white arrows). (**b**) Population data of average transfection rate (>70%) of AAV1-hChR2-YFP in the L5 in both control and cuprizone conditions. (**c**) *Left*, whole-cell current-clamp recording from a PV^+^ interneurons (bottom) overlaid with separate oIPSC recordings from a L5 PN. A 1 sec blue light field illumination (blue bar) produces sustained firing in *PV-Cre AAV1-ChR2* interneurons. *Right*, paired recording of uIPSC in a L5 PN connected with a PV^+^ interneuron revealing a similar brief uIPSC facilitation for the first spikes followed by synaptic depression during a 700 ms train of action potentials. (**d**) *Left*, schematic of whole-cell voltage-clamp recording for mEPSCs in identified parvalbumin interneurons in the *PV-Cre; Ai14* mouse line. mEPSCs were recorded in control mice and mice with 6-weeks cuprizone feeding. (**e, f**) Population data of mEPSC recordings revealed no difference in amplitude nor in the mEPSC frequency. Data show mean ± SEM with open circles individual interneurons. n.s., not significant.

**Fig. S6.**
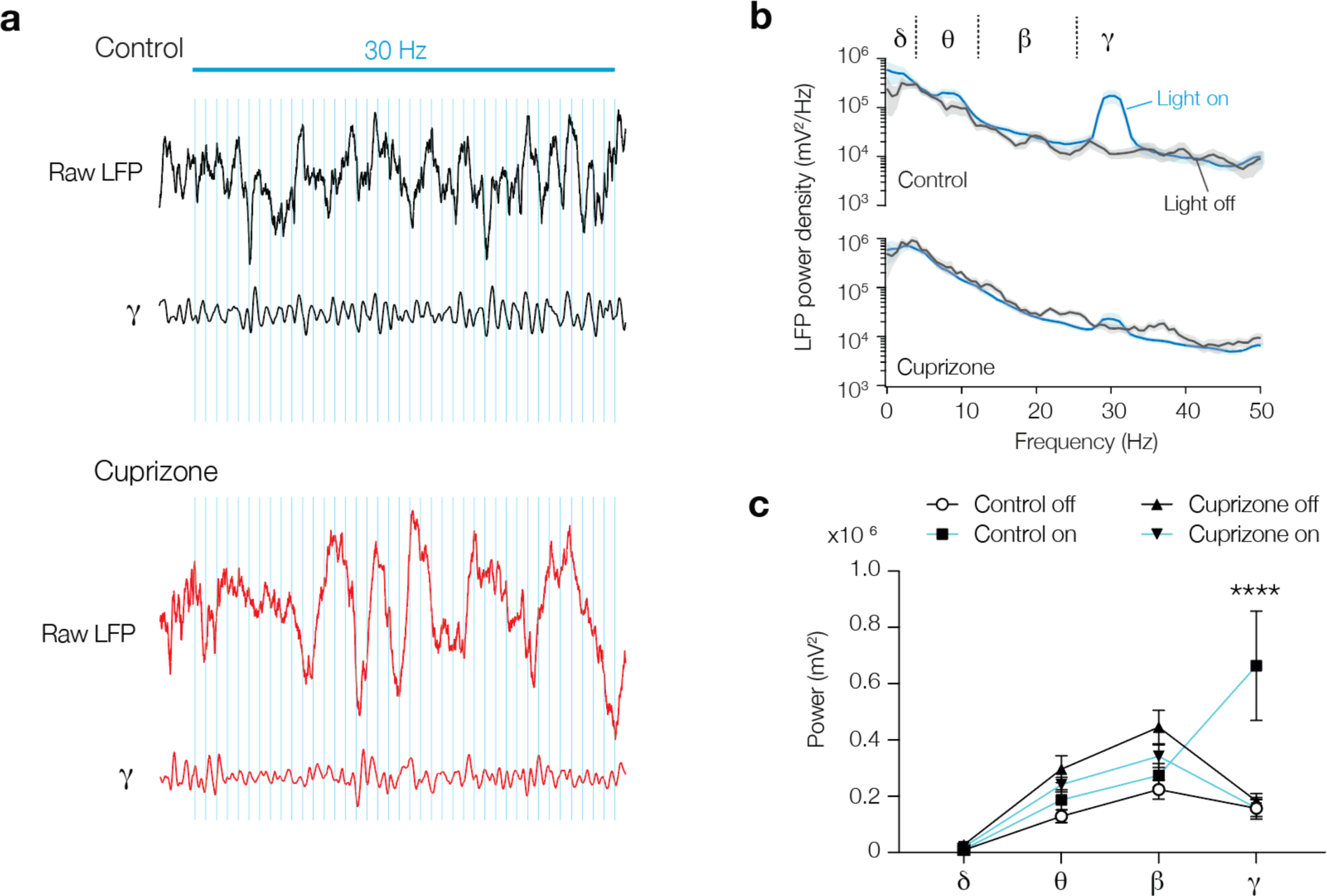
Myelin loss abolishes optogenetically evoked entrainment of γ rhythm. (**a**) Raw LFP and low gamma (γ) 25-40 Hz band-pass filtered trace during 30 Hz blue light stimulation in control (black) and cuprizone (red) mice. Blue light shifted the phase or extended the γ cycle in control mice. (**b**) Averaged power spectral density content of low-γ entrainment during *light on* (blue lines) or *off* (black lines). (**c**) 30 Hz blue light stimulation showed a lack of γ band entrainment in cortex of demyelinated mice in comparison to the significantly increased γ power in control mice. Data show mean ± SEM.

**Fig. S7.**
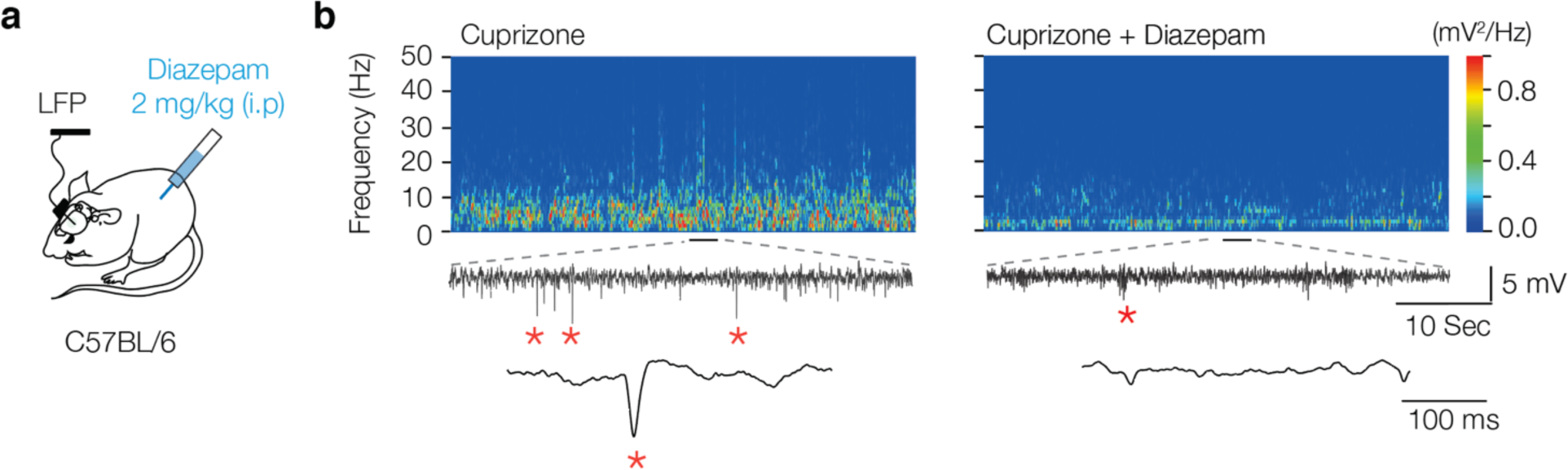
GABA_A_ receptor agonism suppresses interictal epileptiform discharge frequency. (**a**) The GABA_A_ receptor agonist, diazepam, was injected i.p. at 7 weeks of cuprizone treatment, and LFP recordings performed 10 hours post diazepam injection (**b**) Example time frequency plot before (*left*) and after diazepam injection (*right*) showing suppression of interictal epileptiform discharges in cuprizone-treated mice.

**Supplementary Table S1.**
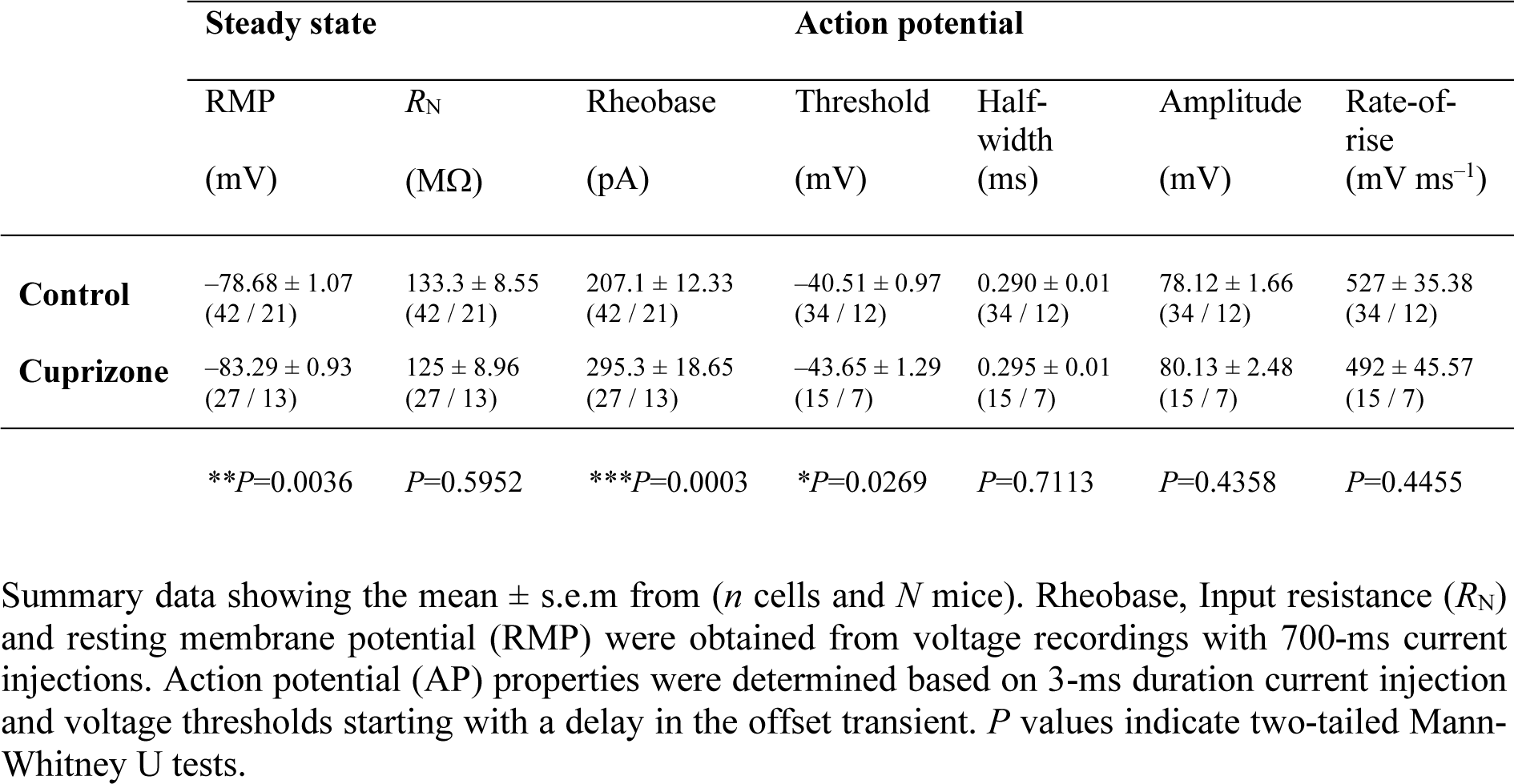
Electrophysiological properties of PV^+^ interneurons.

**Supplementary Table 2. Figure statistics**

See Excel file

**Movie S1. Brain-state dependent interictal spikes**

Example video of ECoG, LFP and behavioral recordings in a 6-weeks cuprizone treated mouse. Raw signals show ECoG from right and left primary somatosensory cortex (S_R_, S_L_, respectively) and LFP from the left layer 5 region (S_L-LFP_). Similar configuration for primary visual cortex (VR, VL and VL-LFP). An additional electrode was connected to the neck muscle recording electromyography (EMG). Note the high EMG activity during awake and moving states. Interictal epileptiform discharges was automatically detected (red asterisks) and only occur during quiet wakefulness.

**Movie S2. Optogenetic activation of myelin-deficient PV^+^ interneurons suppresses interictal spikes**

Example video of ECoG, LFP and behavioral recordings in a 7-weeks cuprizone treated mouse. Raw signals show ECoG from right primary somatosensory cortex (S_R_) and LFP from the layer 5 region (S_R-LFP_). Interictal epileptiform discharges (red asterisks) were suppressed during laser-induced optogenetic activation of PV^+^ interneurons in layer 5.

## References

1. Tremblay, R., Lee, S. & Rudy, B. GABAergic Interneurons in the Neocortex: From Cellular Properties to Circuits. Neuron 91, 260–292 (2016).

2. Hu, H., Gan, J. & Jonas, P. Interneurons. Fast-spiking, parvalbumin^+^ GABAergic interneurons: from cellular design to microcircuit function. Science 345, 1255263 (2014).

3. Bartos, M. et al. Fast synaptic inhibition promotes synchronized gamma oscillations in hippocampal interneuron networks. Proceedings of the National Academy of Sciences of the United States of America 99, 13222–13227 (2002).

4. Gonchar, Y. & Burkhalter, A. Three distinct families of GABAergic neurons in rat visual cortex. Cereb Cortex 7, 347–358 (1997).

5. Tamás, G., Buhl, E. H. & Somogyi, P. Fast IPSPs elicited via multiple synaptic release sites by different types of GABAergic neurone in the cat visual cortex. J Physiology 500, 715–738 (1997).

6. Atallah, B. V., Bruns, W., Carandini, M. & Scanziani, M. Parvalbumin-Expressing Interneurons Linearly Transform Cortical Responses to Visual Stimuli. Neuron 73, 159–170 (2012).

7. Lee, S.-H. et al. Activation of specific interneurons improves V1 feature selectivity and visual perception. Nature 488, 379–383 (2013).

8. Cardin, J. A. et al. Driving fast-spiking cells induces gamma rhythm and controls sensory responses. Nature 459, 663–667 (2009).

9. Zucca, S. et al. An inhibitory gate for state transition in cortex. Elife 6, e26177 (2017).

10. Yang, J.-W. et al. Optogenetic Modulation of a Minor Fraction of Parvalbumin-Positive Interneurons Specifically Affects Spatiotemporal Dynamics of Spontaneous and Sensory-Evoked Activity in Mouse Somatosensory Cortex in Vivo. Cerebral Cortex 27, 5784–5803 (2017).

11. Sommeijer, J.-P. & Levelt, C. N. Synaptotagmin-2 is a reliable marker for parvalbumin positive inhibitory boutons in the mouse visual cortex. PLoS ONE 7, e35323 (2012).

12. Chen, C., Arai, I., Satterfield, R., Young, S. M. & Jonas, P. Synaptotagmin 2 Is the Fast Ca2+ Sensor at a Central Inhibitory Synapse. Cell reports 18, 723–736 (2017).

13. Somogyi, P., Kisvárday, Z. F., Martin, K. A. & Whitteridge, D. Synaptic connections of morphologically identified and physiologically characterized large basket cells in the striate cortex of cat. Neuroscience 10, 261–294 (1983).

14. Thomson, A. M., West, D. C., Hahn, J. & Deuchars, J. Single axon IPSPs elicited in pyramidal cells by three classes of interneurones in slices of rat neocortex. J Physiology 496, 81–102 (1996).

15. Peters, A. & Proskauer, C. C. Smooth or sparsely spined cells with myelinated axons in rat visual cortex. Neuroscience 5, 2079–2092 (1980).

16. Micheva, K. D. et al. A large fraction of neocortical myelin ensheathes axons of local inhibitory neurons. eLife 5, e15784 (2016).

17. Stedehouder, J. et al. Fast-spiking Parvalbumin Interneurons are Frequently Myelinated in the Cerebral Cortex of Mice and Humans. Cerebral Cortex 27, 5001–5013 (2017).

18. Yang, S. M., Michel, K., Jokhi, V., Nedivi, E. & Arlotta, P. Neuron class-specific responses govern adaptive myelin remodeling in the neocortex. Sci New York N Y 370, (2020).

19. Nave, K.-A. & Werner, H. B. Myelination of the nervous system: mechanisms and functions. Annual review of cell and developmental biology 30, 503–533 (2014).

20. Cohen, C. C. H. et al. Saltatory Conduction along Myelinated Axons Involves a Periaxonal Nanocircuit. Cell 180, 311–322.e15 (2020).

21. Schmidt, H. et al. Axonal synapse sorting in medial entorhinal cortex. Nature 549, 469–475 (2017).

22. Micheva, K. D., Kiraly, M., Perez, M. M. & Madison, D. V. Conduction Velocity Along the Local Axons of Parvalbumin Interneurons Correlates With the Degree of Axonal Myelination. Cereb Cortex 31, bhab018- (2021).

23. Barron, T., Saifetiarova, J., Bhat, M. A. & Kim, J. H. Myelination of Purkinje axons is critical for resilient synaptic transmission in the deep cerebellar nucleus. Scientific reports 8, 1022 (2018).

24. Benamer, N., Vidal, M., Balia, M. & Angulo, M. C. Myelination of parvalbumin interneurons shapes the function of cortical sensory inhibitory circuits. Nature Communications 11, 5151 (2020).

25. Veit, J., Hakim, R., Jadi, M. P., Sejnowski, T. J. & Adesnik, H. Cortical gamma band synchronization through somatostatin interneurons. Nat Neurosci 20, 951–959 (2017).

26. Sohal, V. S., Zhang, F., Yizhar, O. & Deisseroth, K. Parvalbumin neurons and gamma rhythms enhance cortical circuit performance. Nature 459, 698–702 (2009).

27. Buzsáki, G. Rhythms of the Brain. (Oxford University Press, 2006). doi:10.1093/acprof:oso/9780195301069.001.0001.

28. Kipp, M., Clarner, T., Dang, J., Copray, S. & Beyer, C. The cuprizone animal model: new insights into an old story. Acta Neuropathologica 118, 723–736 (2009).

29. Hamada, M. S. & Kole, M. H. P. Myelin loss and axonal ion channel adaptations associated with gray matter neuronal hyperexcitability. The Journal of neuroscience 35, 7272–7286 (2015).

30. Clarner, T. et al. Myelin debris regulates inflammatory responses in an experimental demyelination animal model and multiple sclerosis lesions. Glia 60, 1468–1480 (2012).

31. Dubey, M. et al. Seizures and disturbed brain potassium dynamics in the leukodystrophy megalencephalic leukoencephalopathy with subcortical cysts. Annals of Neurology 83, 636–649 (2018).

32. Cohen, I., Navarro, V., Clémenceau, S., Baulac, M. & Miles, R. On the origin of interictal activity in human temporal lobe epilepsy in vitro. Science 298, 1418–1421 (2002).

33. Tóth, K. et al. Hyperexcitability of the network contributes to synchronization processes in the human epileptic neocortex. J Physiology 596, 317–342 (2017).

34. Hoffmann, K., Lindner, M., Gröticke, I., Stangel, M. & Löscher, W. Epileptic seizures and hippocampal damage after cuprizone-induced demyelination in C57BL/6 mice. Experimental neurology 210, 308–321 (2008).

35. Readhead, C. et al. Expression of a myelin basic protein gene in transgenic shiverer mice: Correction of the dysmyelinating phenotype. Cell 48, 703–712 (1987).

36. Chernoff, G. F. Shiverer: an autosomal recessive mutant mouse with myelin deficiency. The Journal of heredity 72, 128 (1981).

37. Brill, J., Mattis, J., Deisseroth, K. & Huguenard, J. R. LSPS/Optogenetics to Improve Synaptic Connectivity Mapping: Unmasking the Role of Basket Cell-Mediated Feedforward Inhibition. eNeuro 3, (2016).

38. Xu, J., Mashimo, T. & Südhof, T. C. Synaptotagmin-1, -2, and -9: Ca(2+) sensors for fast release that specify distinct presynaptic properties in subsets of neurons. 54, 567–581 (2007).

39. Packer, A. M. & Yuste, R. Dense, unspecific connectivity of neocortical parvalbumin-positive interneurons: a canonical microcircuit for inhibition? The Journal of neuroscience 31, 13260– 13271 (2011).

40. Turecek, J., Jackman, S. L. & Regehr, W. G. Synaptic Specializations Support Frequency-Independent Purkinje Cell Output from the Cerebellar Cortex. Cell Reports 17, 3256–3268 (2016).

41. Zhou, J., Lenck-Santini, P., Zhao, Q. & Holmes, G. L. Effect of Interictal Spikes on Single-Cell Firing Patterns in the Hippocampus. Epilepsia 48, 720–731 (2007).

42. Kleen, J. K., Scott, R. C., Holmes, G. L. & Lenck-Santini, P. P. Hippocampal interictal spikes disrupt cognition in rats. Ann Neurol 67, 250–257 (2010).

43. Poulet, J. F. A. & Petersen, C. C. H. Internal brain state regulates membrane potential synchrony in barrel cortex of behaving mice. Nature 454, 881–885 (2008).

44. Gentet, L. J., Avermann, M., Matyas, F., Staiger, J. F. & Petersen, C. C. H. Membrane Potential Dynamics of GABAergic Neurons in the Barrel Cortex of Behaving Mice. Neuron 65, 422–435 (2010).

45. Stedehouder, J. et al. Local axonal morphology guides the topography of interneuron myelination in mouse and human neocortex. eLife 8, (2019).

46. Stedehouder, J. & Kushner, S. A. Myelination of parvalbumin interneurons: a parsimonious locus of pathophysiological convergence in schizophrenia. Molecular psychiatry 22, 4–12 (2017).

47. Miles, R. Variation in strength of inhibitory synapses in the CA3 region of guinea-pig hippocampus in vitro. J Physiology 431, 659–676 (1990).

48. Rossignol, E., Kruglikov, I., Maagdenberg, A. M. J. M. van den, Rudy, B. & Fishell, G. CaV 2.1 ablation in cortical interneurons selectively impairs fast-spiking basket cells and causes generalized seizures. Annals of Neurology 74, 209–222 (2013).

49. Casale, A. E., Foust, A. J., Bal, T. & McCormick, D. A. Cortical Interneuron Subtypes Vary in Their Axonal Action Potential Properties. J Neurosci 35, 15555–15567 (2015).

50. Hu, H. & Jonas, P. A supercritical density of Na(+) channels ensures fast signaling in GABAergic interneuron axons. Nature Neuroscience (2014) doi:10.1038/nn.3678.

51. Lubetzki, C., Sol-Foulon, N. & Desmazieres, A. Nodes of Ranvier during development and repair in the CNS. Nature Reviews Neurology 16, 426–439 (2020).

52. Freeman, S. A. et al. Acceleration of conduction velocity linked to clustering of nodal components precedes myelination. Proceedings of the National Academy of Sciences 112, E321–8 (2015).

53. Micheva, K. D. et al. Distinctive Structural and Molecular Features of Myelinated Inhibitory Axons in Human Neocortex. eNeuro 5, (2018).

54. Edgar, J. M. et al. Early ultrastructural defects of axons and axon-glia junctions in mice lacking expression of Cnp1. Glia 57, 1815–1824 (2009).

55. Fünfschilling, U. et al. Glycolytic oligodendrocytes maintain myelin and long-term axonal integrity. Nature 485, 517–521 (2012).

56. Berg, R. van den, Hoogenraad, C. C. & Hintzen, R. Q. Axonal transport deficits in multiple sclerosis: spiraling into the abyss. Acta Neuropathologica 134, 1–14 (2017).

57. Lindner, M., Fokuhl, J., Linsmeier, F., Trebst, C. & Stangel, M. Chronic toxic demyelination in the central nervous system leads to axonal damage despite remyelination. Neuroscience letters 453, 120–125 (2009).

58. Sorbara, C. D. et al. Pervasive Axonal Transport Deficits in Multiple Sclerosis Models. Neuron 84, 1183–1190 (2014).

59. Chen, Z. et al. Microglial displacement of inhibitory synapses provides neuroprotection in the adult brain. Nature communications 5, 4486 (2014).

60. Ramaglia, V. et al. Complement-associated loss of CA2 inhibitory synapses in the demyelinated hippocampus impairs memory. Acta Neuropathol 1–25 (2021) doi:10.1007/s00401-021-02338-8.

61. Favuzzi, E. et al. GABA-receptive microglia selectively sculpt developing inhibitory circuits. Cell 184, 4048–4063.e32 (2021).

62. Caprariello, A. V. et al. Biochemically altered myelin triggers autoimmune demyelination. Proc National Acad Sci 115, 201721115 (2018).

63. Skripuletz, T. et al. Astrocytes regulate myelin clearance through recruitment of microglia during cuprizone-induced demyelination. Brain 136, 147–167 (2013).

64. Poggi, G. et al. Cortical network dysfunction caused by a subtle defect of myelination. Glia 64, 2025–2040 (2016).

65. Benedict, R. H. B., Amato, M. P., DeLuca, J. & Geurts, J. J. G. Cognitive impairment in multiple sclerosis: clinical management, MRI, and therapeutic avenues. Lancet Neurology 19, 860–871 (2020).

66. Kleen, J. K. et al. Hippocampal interictal epileptiform activity disrupts cognition in humans. Neurology 81, 18–24 (2013).

67. Henin, S. et al. Spatiotemporal dynamics between interictal epileptiform discharges and ripples during associative memory processing. Brain (2021) doi:10.1093/brain/awab044.

68. Lam, A. D. et al. Silent hippocampal seizures and spikes identified by foramen ovale electrodes in Alzheimer’s disease. Nat Med 23, 678–680 (2017).

69. Chen, J.-F. et al. Enhancing myelin renewal reverses cognitive dysfunction in a murine model of Alzheimer’s disease. Neuron 109, 2292–2307.e5 (2021).

70. Drenthen, G. S. et al. On the merits of non-invasive myelin imaging in epilepsy, a literature review. J Neurosci Meth 338, 108687 (2020).

71. Zoupi, L. et al. Selective vulnerability of inhibitory networks in multiple sclerosis. Acta Neuropathol 141, 415–429 (2021).

72. Tewarie, P. et al. Disruption of structural and functional networks in long-standing multiple sclerosis. Human brain mapping 35, 5946–5961 (2014).

73. Schoonheim, M. M. et al. Functional connectivity changes in multiple sclerosis patients: a graph analytical study of MEG resting state data. Human brain mapping 34, 52–61 (2013).

74. Cawley, N. et al. Reduced gamma-aminobutyric acid concentration is associated with physical disability in progressive multiple sclerosis. Brain 138, 2584–2595 (2015).

75. Gao, F. et al. Altered hippocampal GABA and glutamate levels and uncoupling from functional connectivity in multiple sclerosis. Hippocampus 28, 813–823 (2018).

